# The Loss of the H1.4 Linker Histone Impacts Nascent Transcription and Chromatin Accessibility

**DOI:** 10.1101/2023.05.14.540702

**Authors:** Nolan G. Gokey, James M. Ward, Eric J. Milliman, Leesa J. Deterding, Kevin W. Trotter, Trevor K. Archer

**Author notes:** Corresponding Author: Trevor K. Archer, PhD, P.O Box 12233, Research Triangle Park, NC 27709.

## Abstract

The Chromatosome superstructure, comprised of core histone containing nucleosomes and linker histones, act in concert as physical barriers to genetic material in the mammalian nucleus to trans-acting factors. Appropriate arrangement, composition, and post-translational modification of the chromatosome is highly regulated and necessary for appropriate gene expression. These proteins act to radically condense the genetic material and linker H1 histone is essential for the further condensation of the chromatin fiber. However, the regulatory role of H1 in gene expression and chromatin organization is complicated by cell type specific expression and compensation of multiple H1 variants. Leveraging the UL3 osteosarcoma cell line which displays biased expression of H1 variants, and CRISPR/Cas9, we generated H1.4-deficient clones. Loss of H1.4 results in consistent changes to chromatin accessibility concomitant with changes to histone tail modifications, as well as a set of differentially expressed genes shared among ΔH1.4 genetic clones. We identified immune and inflammation immediate early genes as enriched in differentially expressed genes, skewed towards AP-1 regulated targets. Our data show that H1.4 is critical for the regulation of stress response pathways.

**Key Points for NAR:** (3 bullet points summarizing the manuscript’s contribution to the field)

- H1.4 is essential for appropriate expression of over 6,000 nascent transcripts in UL3 cells.
- Loss of H1.4 results in widespread changes in chromatin accessibility at enhancers and transcribed regions as well as heterochromatin and quiescent chromatin.
- Immediate early genes, and especially AP-1 family members, are highly sensitive to H1.4 loss and their binding sites coincide with losses in chromatin accessibility

## INTRODUCTION

Chromatin modulates gene activity by regulating access to genomic material through the location and post-translation modification of histones (1–5). Placement, eviction, and posttranslational modification of core nucleosomal proteins (H2, H3, H4) play a pivotal role in the regulation of gene activity (6–10). However, less is understood about the role of the linker histone, H1, which binds at the nucleosomal-DNA dyad and acts to bridge linker DNA and form the chromatosome superstructure (11–15). While the nucleosome residence time has been determined to be >130min (16), H1 turnover on chromatin occurs on the scale of <4 minutes and occurs nearly continuously (17,18). Consistent with this observation, arrangement, and modification of H1 is highly dynamic and highly responsive to cellular stimuli. H1 aids chromosomal compaction, stabilize nucleosomes on DNA, and prevent their dislocation (19). Displacement of H1 along with the maintenance of nucleosomal localization has been shown to alleviate the transcriptional repression of chromatin (20).

H1’s primary function with regards to gene regulation is thought to be repressive through compaction of chromatin and stabilization of the core nucleosome (16–19). Work in hepatocytes has shown that mere displacement of H1 by pioneer transcription factor FoxA is enough to induce gene activation, even in the presence of stable nucleosomes (20). We and others have shown histone H1 to play a mechanistic and regulatory role regulating the Mouse Mammary Tumor Virus (MMTV) promoter in response to corticosteroid treatment, further highlighting H1’s important role in gene expression control and changes to chromatin architecture with response to stimuli (21–28). Beyond a gene-specific role of H1 in controlling chromatin occupancy, H1 also plays an architectural role and as such has domain-level roles in organizing each chromosome and condensing the nucleus (11,29,30).

The H1 family has 11 members and consists of two families; the variants encoded as a gene cluster (H1.1, H1.2, H1.3, H1.4, H1.5 and H1.6) and the orphan gene variants (H1.0, H1.7, H1.8, H1.9 and H1.10). Examination of the specific role of each linker histone variant by siRNA and genetic engineering has been hampered by the existence of compensatory effects between variants. Moreover, whole body-ablation in mice of a single H1 variant resulted in little to no phenotype until multiple variants were deleted at which point the animals died early in life (31). Losses of single linker histone variants or even pairs of variants within the H1 gene cluster resulted in normal development (32,33). Conversely, each of the six human cluster 1 linker H1 variants, while structurally similar, has unique roles in nuclear organization due to both the unique primary sequences of the variants as well as the complex mix of variant-specific post-translational modifications often decorating the linker histone (34).

Herein we use whole genome approaches, genetic engineering, and intrinsic factors to investigate the contributions of variant H1.4 to global gene and chromatin regulation. To date, H1.4 is the only H1 variant linked to clinical human disease, with mutations in the C-terminal domain of H1-4 (formerly HIST1H1E) being linked to Overgrowth and Intellectual Disability (39). We take advantage of the intrinsic H1 variant expression characteristics of U-2 OS human osteosarcoma cancer cells (35,36) which have been reported to express predominantly variant H1.4 (23,24,37). Using a CRISPR/Cas9 genome editing strategy, we generated multiple UL3 clonal lines with disruption of H1.4 gene expression. Using newer nonbiased next-generation sequencing techniques, we perform genome-wide assessment of the effect of H1.4 absence on the arrangement of chromatin, on core histone modifications, as well as on total and nascent gene expression. Our findings further highlight the importance of histone H1.4 in both gene regulation and chromatin organization, while suggesting a specific role for the H1.4 variant in cell types that respond to environmental stressors.

## METHODS AND MATERIALS

### CRISPR Inactivation of H1-4 Locus

The U-2O-S UL3 subclone used was generated and maintained as previously (37) described with the following changes. UL3 cells were maintained in DMEM high glucose (Gibco 11965-092) supplemented with 4mM glutamine, 10% fetal bovine serum (Atlanta Biologicals S11150) and 1% penicillin-streptomycin (Sigma P0781). Cells were maintained in a 5% CO_2_ environment at 37°C and never allowed to exceed 80% confluency. 0.25% Trypsin-EDTA (Gibco 25200) was used following the manufacturer’s instructions for all passaging and cell collection steps. CRISPR/Cas9 nickase gene editing was performed using vector pX335-U6-Chimeric BB-CBh-hSpCas9n(D10A) (Addgene) modified by standard restriction enzyme cloning (BbsI) to contain specific guide sequences targeting the N-terminal portion of the globular domain of H1-4 (see figure S1). Transfections were performed following the manufacturer’s instructions for Lipofectamine 2000 (Life Technologies 11668030). Clonal cell lines were isolated from colonies formed from single cell plating. Genetically distinct cell lines were derived using independent gDNA; guide1 for ΔH1-A, ΔH1-D and ΔH1-E and guide 2 for ΔH1-B and ΔH1-C ΔH1.4 CRISPR clones (oligo sequences Supplemental Table 1). Examination of the CRISPR editing events in individual clones was performed using Sanger sequencing and deconvolution of potential alternative allele sequences was performed using PolyPeakParser (38). Five individual clones were used for subsequent experiments.

### Acid extraction of nuclear proteins and tandem mass spectrometry

An equal number of cells from each H1.4 variant cell line was wash with PBS, resuspended in 5% perchloric acid (244252 Sigma) and kept on ice for 2h with occasional inversion to mix. Samples were centrifuged (13,000xg, 10min, 4°C), and the supernatant transferred to a new tube for precipitation of total with 30% trichloroacetic acid (T4885 Sigma) overnight at −20°C. Precipitated proteins were washed with ice cold acetone and allowed to air dry on ice. Proteins were resuspended in cold water and the protein concentration determined by Bradford assay. Ammonium bicarbonate was added to the samples to a concentration of 0.1ug/uL and each sample was digested with trypsin (1ug/uL) overnight at 37°C. The digests were then stored at −80°C until analysis by mass spectrometry (MS). Protein digests were analyzed by LC/MS on either a Premier Q-Tof mass spectrometer (Waters Corp.) or a Q Exactive Plus mass spectrometer (ThermoFisher Scientific) and a UPLC system equipped with a 75µm x 150µm BEH dC18 column (1.8µm particle, Waters Corporation) and a C18 trapping column (180µm × 20µm) with 5mm particle size at a flow rate of 450nL/min. The trapping column was positioned in-line of the analytical column and upstream of a micro-tee union which was used both as a vent for trapping and as a liquid junction. Trapping was performed using the initial solvent composition. 5µL of digested sample was injected onto the column. Peptides were eluted by using a linear gradient from 97% solvent A (0.1% formic acid in water (v/v)) and 3% solvent B (0.1% formic acid in acetonitrile (v/v)) to 35% solvent B over 70min then ramped to 85% solvent B for 2min. For the mass spectrometry, a top-ten data dependent acquisition method was employed with a dynamic exclusion time of 15sec and an exclusion of +1 charge states. Mass calibration was performed before data acquisition. Quantitation was performed by spiking into each sample a tryptic digest of yeast alcohol dehydrogenase before analysis. Raw mass spectra data were loaded in batch to the Proteome Discoverer software (ThermoFisher Scientific) and analyzed in comparison to the human UniProt library of proteins. Normalized Abundance values for all observed peptides were log2-transformed and examined by MA plot, and two outliers with mean absolute deviation (MAD) scores above 2 were eliminated. Technical and biological replicates were assessed for clonal agreement (Supplemental Figure 1F). Clustered and orphan H1 variant data was condensed to peptides, to proteins, and finally to cell lines (Figures 1B). A threshold of a max group mean of log2 9.5 or 500 Normalized Abundance was applied before performing a limma moderated t-test for differential analysis. 9 biological UL3 replicates and 6 biological replicates from each ΔH1-A and ΔH1-B clones were analyzed, 3 biological replicates for ΔH1-E and 2 biological replicates for ΔH1-C and ΔH1-D. Technical quadruplicates of parental cells and clonal cell lines were also performed, for a total of 105 mass spectrometry samples analyzed.

**Figure 1.**
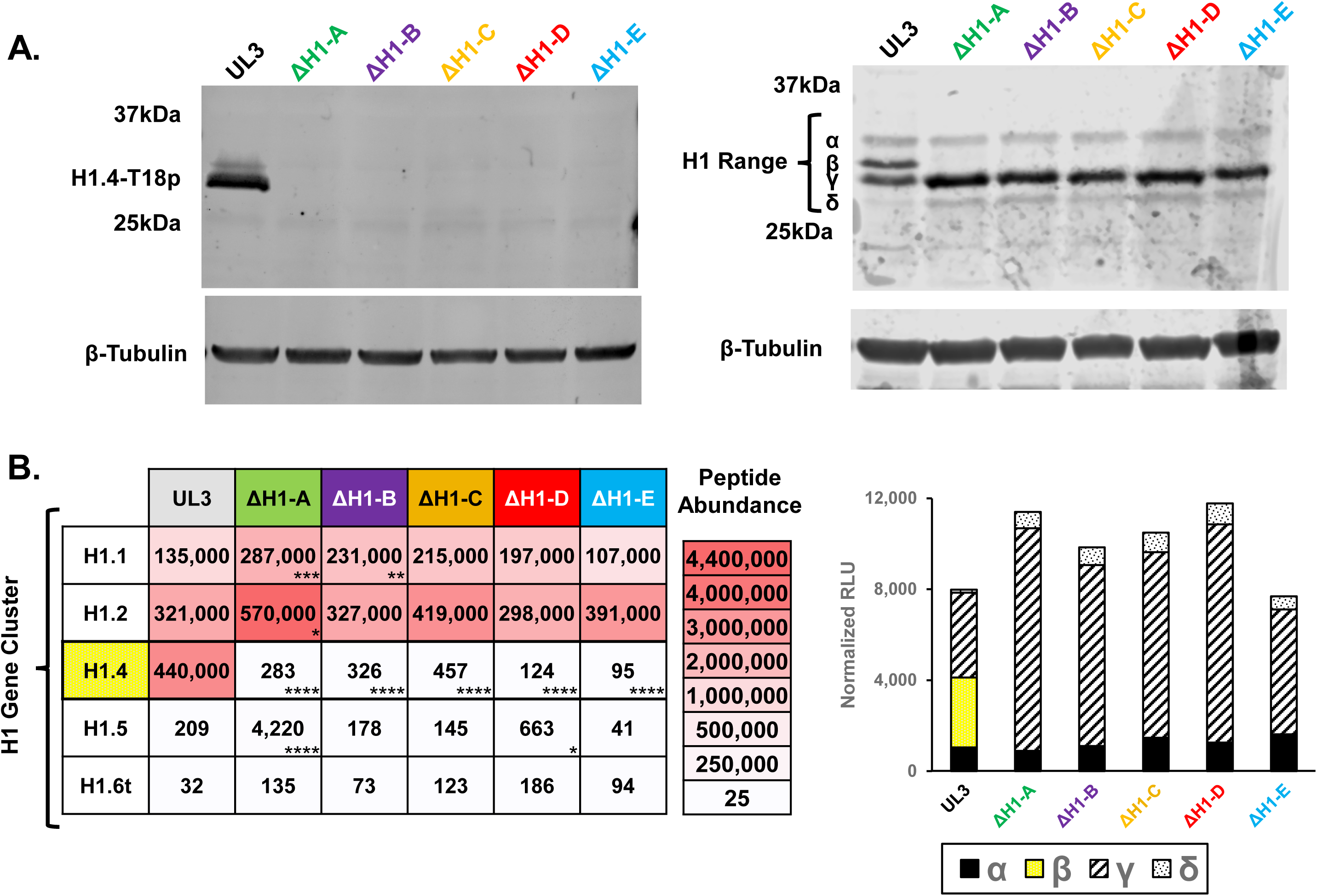
Verification of the loss of H1.4 protein and changes in the overall H1 variant pool in ΔH1.4 clones as compared to UL3 parental cells. **A.** Western Blot results of protein isolates from parental UL3 cells and five ΔH1.4 CRISPR/Cas9 clones. Specific visualization of H1.4 protein was possible using an antibody specific to the dominant H1.4 post-translational modification, H1.4pT18 (left). Visualization of all H1 variants was possible using a pan-H1 antibody (right) and bands are putative H1 bands are labeled with the Greek letters α, β, γ, δ with H1.4 being represented by β. For both H1 Western blots, tubulin was used as a protein loading control (bottom panels). Densitometry for each putative H1 band from the pan-H1 blot is shown below. **B.** Heatmap representation of MS/MS spectral count data for the H1 gene cluster variants (rows) from analyses from parental UL3 cells and H1.4 knockout lines (columns). Adjusted p-value thresholds for variant changes relative to UL3 cells are indicated with asterisks (*≤0.05, **≤0.01, ***≤0.001 and ****≤0.00001).

### Western Blot and Quantitative Gel Staining

Cells were collected with 0.25%Trypsin-EDTA, washed with cold PBS and counted. Cell lysis and denaturing of proteins was performed using Laemmli sample buffer (Bio-Rad 1610737), 100mM DTT, and boiling. Denatured protein lysates (3X10^5^ cells) and protein ladders (Bio-Rad 1610374) were resolved by SDS-PAGE on a 10% Tris-HCl Criterion acrylamide gel. SDS-PAGE western blot gels were transferred to nitrocellulose membranes using a Bio-Rad Trans-Blot Turbo and standard transfer conditions. Membranes were stained with Ponceau S solution (ThermoFisher A40000279), washed with water, and incubated blocking solution (5% milk in TBS) for a minimum of 1 hour with gentle rocking. Membranes were probed with primary antibodies diluted in blocking solution for 1-2 hours at room temperature and with gentle rocking, followed by washing with TBS plus 1% Tween-20. Membranes were probed with secondary antibodies diluted in blocking solution for 1-2 hours at room temperature and with gentle rocking, followed by washing with TBS. Blots were imaged using a Li-Cor Odyssey CLx and results processed and quantified with Image Studio v5.2. Primary antibodies used: β-tubulin (DHSB E7), GAPDH (DSHB 2G7), H1.0 (ab125027 Abcam), H1.1 (HPA043753 Sigma), H1.2 (ab4086 Abcam), H1.4-T18p (37). Secondary antibodies used: IR680 and IR800 antibodies (LiCor 926-68074, 926-68072, 926-68073, 926-32212, 926-32213).

### ATAC-seq

Tagmentation was performed using 50,000 nuclei (isolated as previously described, 45) in 25ul of 1X tagmentation buffer with 3.75ul of Tn5 enzyme for 30 minutes. ATAC-seq library generation and sequencing was performed as previously described (39–41). ATAC-seq reads were deduplicated (Picard) and mapped to hg19 using Bowtie (42). Read pairs were converted to fragments by filling in coverage between the pairs. Open chromatin ATAC-seq peaks were defined using coverage regions and were called in each biological replicate using MACS2 using the parameters --nomodel --nolambda --keep-dup all --slocal 10000 -q 0.005 (39). The peaks from ATAC-seq biological replicates were merged (bedtools) and regions with overlap of structural regions of the genome erroneously captured by ATAC-seq (e.g. centromeres, repeats) were masked (43). Each Differentially Accessible Chromatin (DAC) was annotated relative to the hg19 genome by overlap with annotated genic regions as defined: promoter proximal (- 2000 to +200bp from the annotated transcription start site (TSS), exonic, or intronic. DACs lacking overlap with these genic regions were labeled extragenic. The total genic space was defined by using only the gene proximal promoter regions with at least one ATAC-seq peak. The total number of ATAC-seq peaks in the two knockout lines and parental line (Supplemental Figure 2B, 237,458 peaks) were called, peaks merged into one set, and peaks with fewer than 16 reads were removed from downstream analysis. This set of 156,073 combined ATAC-seq peaks represents all genome-wide sites with accessible chromatin. Peaks were tested for differential accessibility with limma-voom (44), applying at least 1.5-fold change with Benjamini-Hochberg adjusted p-value less than 0.05. The steps were implemented in R package slicejam (https://github.com/jmw86069/slicejam), which provides the following: visual data quality control, signal normalization and optional batch adjustment, visual review of normalization, differential analysis, visualization of DAC hits, and output with annotation of DACs. Before unifying replicates, a heatmap was made of batch adjusted ATAC-seq data at the 20,653 called DACs (Supplemental Figure 3C). For data visualization, replicates were depth normalized and merged to form a single union replicate coverage file. In the 32 cases where a DAC had opposing directions of change in the two lines, the grouping observed in the ΔH1-A line was adopted. Differential heatmaps were produced with ComplexHeatmap (45), genome coverage heatmaps were produced with R packages EnrichedHeatmap (46), and coverjam (https://github.com/jmw86069/coverjam). Larger peaks were sliced into 1kb pieces for independent statistical testing of variable sized peaks (Supplemental Figure 2A). This representative set of accessible peaks was used to call DACs in UL3 cells verses ΔH1.4 cell lines (>1.5-fold, padj <0.05, Supplemental Figure 3A). Epilogos chromatin state scores in 200-base bins were used to define the chromatin state with highest score in each bin (47). The 15 Epilogos chromatin states were also grouped into six state groups: TSS, enhancer, transcribed, heterochromatin, polycomb, and quiescent. Each DAC was annotated using the first overlapping Epilogos chromatin state in the order listed, to augment genic context (promoter, exon, intron, extragenic) for each chromatin region. Log-linear analyses were used to analyze the relationship between each subset of ATAC-seq peaks, genic regions, and Epilogos chromatin states. Frequencies were analyzed using R packages MASS and vcdExtra, and significant changes were defined by a Pearson chi-squared residual > 2. Pearson residuals 2 and 4 represent approximately ɑ = 0.05, and ɑ = 0.0001 (48).

### ChIP-seq

Chromatin immunoprecipitation (ChIP) and next generation sequencing of ChIP-seq libraries was performed as described in (49). Briefly, adherent cells were cross-linked with 1% formaldehyde for 10min at 37°C and quenched with glycine. Cells were collected by scraping, washed with PBS, and centrifuged. Cells were washed with hypotonic buffer, resuspended in shearing buffer, and subjected to gentle sonication using a Bioruptor (Diagenode) at 4°C to generate ChIP material. ChIP material fragment size was evaluated by agarose gel and immunoprecipitations performed as previously described (49). 1ug of antibody was used per 1ug of crosslinked ChIP material in immunoprecipitation reactions. Immune complexes captured with protein A/G Dynabeads (10015D, Invitrogen) and subjected to wash and elution buffers. Eluted DNA fragments were further purified using column purification prior to library preparation using Swift DNA (406162642 Swift/IDT) or Illumina Nextera (20018704 Illumina) library preparation kits. 2 biological replicates were performed for all ChIP-seq experiments and compared to negative control immunoprecipitations (IgG) and input data. Antibodies used for immunoprecipitation include anti-H3K27ac Abcam ab4729, anti-H3K4me1 Abcam ab8895.

### RNA-seq, 4sU- seq, and RT-qPCR

Total RNA was isolated from cells with Trizol (15596018 Ambion) followed by a further acid phenol:chloroform (AM9720 Ambion) and the nucleic acid ethanol precipitated. Prior to RNA isolation, cells were metabolically labelled by exposure to media containing 700uM 4-thio-uracil for 9min at 37°C for 4sU-seq experiments as in (50–54) as well as the advanced biotinylating reagent (53). Samples were treated with DNase I (18068015 Invitrogen), acid phenol extracted, and ethanol precipitated. RNA was linearized at 65°C for 2min prior to concentration assessment and 300ug of RNA was mixed with 10X biotinylation buffer and 100ng of Biotin-XX-MTSEA (90066-1 Biotonium) per ug of RNA. Biotinylation reactions proceeded for 2h at room temperature and with gentle rotation followed by acid phenol extraction and precipitation. Biotinylated RNAs were captured using streptavidin magnetic beads (Promega), washed with salt buffers, and eluted with DTT containing elution buffer. RT-qPCR (methods below) was performed to assess the enrichment quality of nascent transcripts as compared to mature transcripts. RNA-seq Library preparation and sequencing was performed by Expression Analysis/Quintiles (Research Triangle Park, NC). 4sU-seq libraries were prepared using the TruSeq Stranded small RNA seq (R S-200-0012). Total RNA-seq was performed using the RiboZero TruSeq Stranded total RNA sequencing kit (Illumina). All RNA-seq experiments were performed in at least biological duplicate.

Cutadapt (55) was used to remove low quality reads and remove Illumina adapter sequence. Read pairs were mapped to the hg19 genome build with STAR (56), and featureCounts (57) (Bioconductor) and DESeq2 (58) were used to determine changes in gene expression for each transcript with a threshold of a fold change of |1.5| and an Benjamini-Hochberg adjusted p-value less than 0.05. For visualization on UCSC, the depth normalized libraries were merged between biological replicates and bigwig tracks rendered. After identification of DEGs in the 4sU-seq experiments, each set of DEGs were loaded in Ingenuity Pathway Analysis (Qiagen) and examined for shared dysregulation of pathways in the two knockout lines relative to the parental UL3 cells. For data filtering, gene sets were required to have an adjusted p-value enrichment score below 0.05 and involve three or more significant genes per set. Following DESeq2 identification of either shared or cell-line specific changes in gene expression, RT-qPCR was used to validate chosen DEGs in biological triplicate RNA samples independent of those samples sequenced. Full-length RNA-seq data from the six knockout lines were analyzed using Salmon (59), and fold-cutoffs were adjusted to log2fold change and a adjusted p-value of <0.01. Pathway analysis was performed using DAVID (60,61), and pathway results were compared across H1 knockout cell lines using R package multienrichjam (https://github.com/jmw86069/multienrichjam). Briefly, pathways were hierarchically clustered using the gene-pathway incidence matrix, forming four pathway clusters, subsequently visualized as a Concept Network to display gene-pathway relationships (62).

RNA for qPCR was purified from cells as described above. cDNA was generated using a mix of poly-T and random hexamer oligos and the SuperScript II Reverse Transcriptase system (18064022 Invitrogen). RT-qPCR reactions were performed using SsoAdvanced Universal SYBR Green Supermix (1725275 Bio-Rad), the CFX384 Real-Time System (Bio-Rad), and data analyzed using the Bio-Rad CFX Maestro v1.1 software. All real-time experiments were run in technical quadruplicate and in at least biological triplicate and cycle threshold values were normalized to the geometric mean of four housekeeping genes (PolR3GL, TBP, TubA1B and GAPDH). All primers (Supplemental Table 1) were obtained from Integrated DNA Technologies (Coralville, Iowa).

## RESULTS

### Generation of Linker Histone H1.4 Deficient Osteosarcoma Cells

The Human osteosarcoma U-2O-S cell line H1 gene cluster pool is dominated by the H1.4 linker variant at the protein level (23,24). First, we evaluated the transcript and protein levels of all H1 variants in our UL3 cell line (Supplemental Figure 1A, grey bars), which are U-2O-S cells which we previously generated to contain an MMTV expression cassette and found both H1.0 (orphan) and H1.4 (gene cluster) to be highly expressed. Using the CRISPR/Cas9 nickase system, we generated small indel mutations in a portion of the H1-4 gene coding for amino acids 30-38 the unstructured N-terminal domain (Supplemental Figure 1C). Clones generated single-cell derived colonies and screened for loss of H1.4 using an antibody specific for H1.4-phospho-tyrosine-18, the predominant post-translational modification on H1.4 (23,24). Five UL3 ΔH1.4 clones with verified absence of H1.4 protein expression were selected for use in downstream experiments, designated as clones A-E (Figure 1A). The calculated molecular weight of H1.4 is 22kDa but migrates considerably larger at ∼33kDa likely due to considerable post-translational modifications (63,64). These clones were further verified using Sanger sequencing to confirm that each clone harbors a unique indel mutation and likely arose from an individual CRISPR event (Supplemental Figure 1D). As expected, all five clones harbor indel mutations located at the junction between the N-terminal disordered region and the globular domain, resulting in a frameshift, early termination and loss of the functional globular domain shared by all H1 linker proteins (Supplemental Figure 1E). While we did not observe an increase in necrotic death as previously reported (27), we did observe that the H1.4-deficient cells grew at a marginally slower rate and as such had to be plated more densely when paired with UL3 parental cells to obtain matched cell counts for experimental isolations (data not shown).

### Ablation of H1.4 protein results in compensation by H1.0, H1.1 and H1.2

Using RT-qPCR and mass spectrometry (MS) with extensive replicates for depth confidence (n=105), we assessed the levels of H1 variant expression in the parental UL3 parental and ΔH1.4 cell lines. Consistent with the observed transcript levels (Supplemental Figure 1A), the UL3 parental line was observed to exhibit little to no peptide signal from H1.5 or H1.6t (Figure 1B) among the H1 gene cluster variants while the orphan H1 variants are dominated by H1.0 (Supplemental Figure 1F).

Across the five ΔH1.4 CRISPR/Cas9 clones, the H1 compensatory profile relative to UL3 cells was remarkably similar at both the transcript and protein level with the H1.1 and H1.2 cluster variants being upregulated, however in some cells this upregulation did not reach statistical significance. We also observed a statistically significant activation of H1.5 in two lines (ΔH1-A and ΔH1-D), a variant not observed in the parental line. Compensatory expression by orphan H1.0 histone has been reported when H1.4 expression was reduced (27) and we also observe upregulation of H1.0 at both the transcript and protein level (Supplemental Figure 1A, 1B). While this variant is more typically expressed in most-mitotic cells or in cells which have exited the cell cycle (65), there is also considerably evidence that H1.0 is highly expressed in rapidly dividing cell cultures (66,67). We saw a more modest level of UL3 expression of orphan variant H1.10 as well as some compensatory upregulation of this orphan in the ΔH1.4 clones (Supplementary Figure 1G).

Changes in the H1 variant protein pool were validated by Western blot using an antibody which recognizes multiple H1 variants (Figure 1A). While H1 variants migrate with different patterns in different cells, the β-band corresponds to the size of H1.4 and the γ-band corresponds to the size of H1.0 (Supplemental Figure 1B), and these bands match the observations from the MS experiments. For further verification of variant protein abundance, Western blots were performed using H1 variant specific antibodies against H1.0, H1.1 and H1.2 (Supplemental Figure 1B) all of which supported the MS results. While commercially available antibodies for H1.5 and H1.6 exist, in the UL3 system these antibodies were found to cross-react with the exponentially higher levels of H1.0, H1.1 and H1.2 relative to the very low levels of H1.5 and H1.6 observed in the cells (data not shown). No unique peptides were observed for variants H1.3, H1.7, H1.8 and H1.9 is an expressed pseudogene in humans (data not shown, see GEO and PRIDE data archive).

### H1.4 loss results in reproducible dysregulation of inflammatory and integrin signaling

Once the five H1.4-knockout lines had been established, and their changes to the H1 variant pool found to be similar, we investigated which genes were responsive to the loss of H1.4. We performed random hexamer RiboZero RNA-seq (Illumina) to enrich for full length transcript coverage. Following quantification of transcript changes relative to UL3 cells, we identified differentially expressed genes (DEGs) in each H1.4 knockout line which are represented as Volcano plots (Figure 2A). Across the five clones, we did not observe a global bias towards upregulated to downregulated expression (Figure 2A).

**Figure 2.**
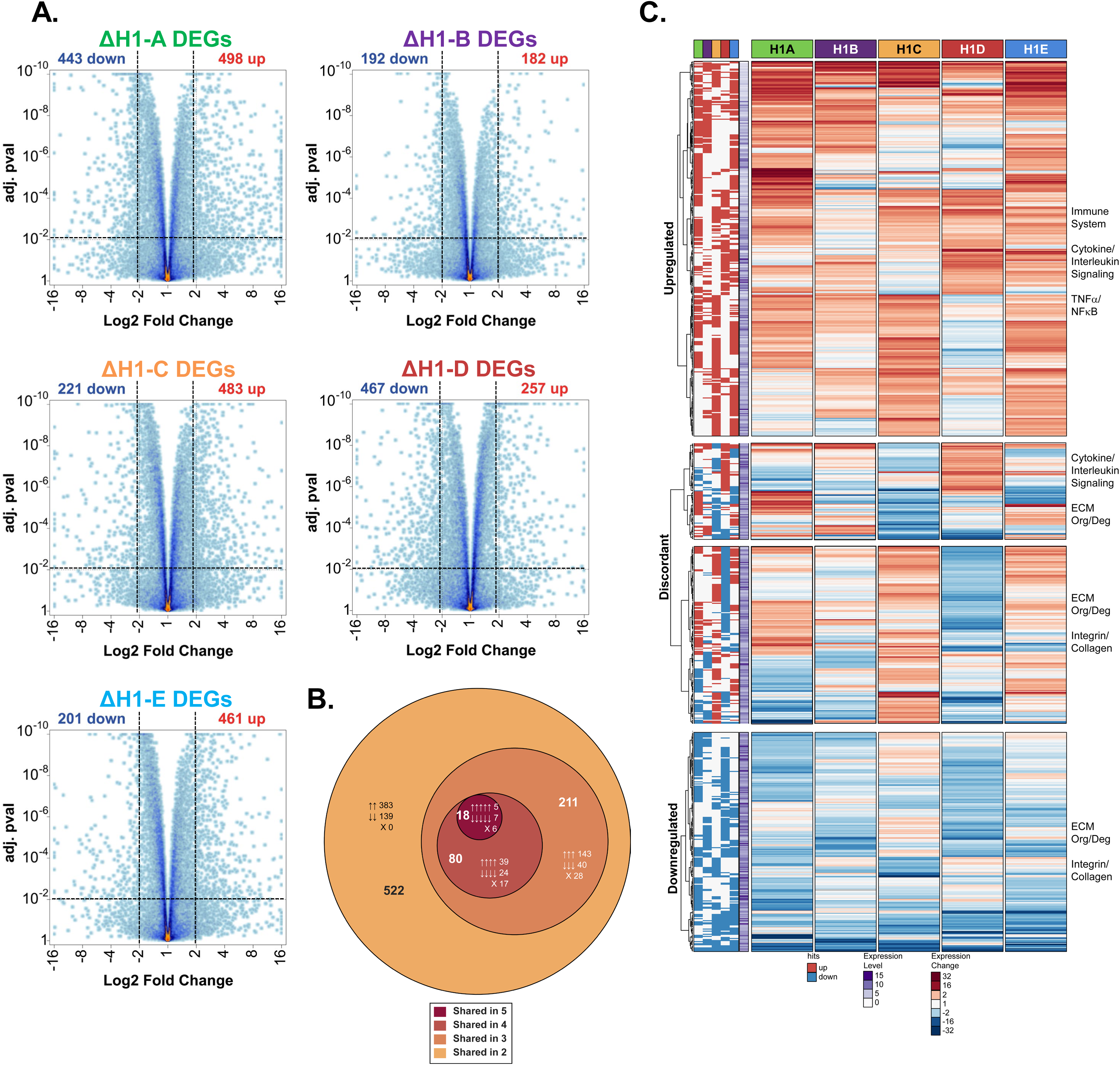
Loss of H1.4 results in a reproducible altered gene expression profile. **A.** RNA-seq DEGs are depicted as volcano plots for each H1.4-knockout cell line (ΔH1-A through D) relative to UL3 cells. The total number of upregulated (red) and downregulated (blue) genes displayed above each quadrant of the volcano plot. Horizontal dashed lines indicate the adjusted p-value threshold (0.01), and vertical dashed lines indicate the 2-fold change threshold for significance. **B.** Proportional Venn diagram showing DEGs shared across the five ΔH1.4 clones. The number of DEGs is shown in each ring as well as the indication of if the gene shares directionality indicated by arrows, with a gene that is dysregulated indicated by an X. **C.** A heatmap of 831 genes called as DEGs in at least 2 found to be dysregulated in at least two ΔH1.4 clones. Binary directional changes are summarized (left columns) adjacent to mean expression data (right columns) for the biological replicate data obtained from each ΔH1.4 clone, and clustered by direction of change. Relevant enriched pathways (DAVID) are indicated for each cluster (right) and the relative expression of each gene in UL3 cells is depicted in the purple vertical stripe. DAVID pval<0.05.

Despite the loss of this critical chromatin factor, most genes maintained their level of processed mRNA in the ΔH1.4 clones. Each ΔH1.4 clone also exhibited a unique set of clone-specific DEGs, with 831 total genes being called DEGs between at least 2 ΔH1.4 clones (Figure 2B). Using a step-down analysis, we observed that 522 of these shared DEGs also have 100% directional agreement (Figure 2B), with the trend continuing (211 DEGs in three lines with 87% concordance, 80 DEGs in four lines with 79% concordance, and the 18 in five lines with 67% concordance). A random sort of DEGs produces only 3% agreement across the five ΔH1.4 clones.

To determine if there was a shared pattern of gene expression in cells lacking H1.4, we performed hierarchical clustering on the 831 shared DEGs identified by RNA-seq and using DAVID (61,68) identified four clusters of expression types in our five knockout lines (Figure 2C). We observed a set of broadly upregulated genes which were identified as belonging to immune system responses, cytokine and interleukin signaling as well as specific pathways such as TNFα and NF-κB. A second cluster of gene responses was identified as downregulated, and these genes play roles in the organization and degradation of the extracellular matrix, collagen formation, as well as integrin signaling (Figure 2C, bottom, pval<0.05). Two other clusters were found to be discordantly dysregulated across the five knockout lines; however, the same pathways were identified within these clusters as those identified as up and downregulated (Figure 2C, middle, pval<0.05). RNA-seq findings were validated with RT-qPCR on a small subset of the DEGs including both repressed and activated genes (Supplemental Figure 7).

### Loss of H1.4 results in Changes to Accessible Chromatin

After establishing that the five ΔH1.4 CRISPR/Cas9 clones share the same H1 compensatory profile at the protein level, and the gene expression changes are shared across the clones, we selected lines ΔH1-A and ΔH1-B for further investigation of genomic changes. To examine the role H1.4 plays in chromatin assembly and organization, we used ATAC-seq to identify changes in chromatin accessibility. In the three lines, we identified 156,073 peaks of open chromatin. (Supplemental Figure 3B). and within these identified peaks which change relative to the UL3 cell line and termed these peaks Differentially Accessible Chromatin (DAC). In the ΔH1-A line we observed 14,924 DACs and 7,964 DACs in the ΔH1-B line for a total of 20,621 DACs (Figure 3A). A slight bias toward a loss of accessibility was observed, with 55% of the DACs in the ΔH1-A line showing a reduction in accessibility and 70% of the ΔH1-B DACs having decreased accessibility (Figure 3A). We were surprised to find the direction of change in chromatin accessibility to be highly concordant between our two H1.4-deficient lines, with 86% of the DACs agreeing in their directional change, or 17,746 different peaks (Figure 3D) a ratio much higher than one would expect by chance. This is best visualized in a heatmap of all 20,653 DACs, showing agreement in direction of change at most sites of differential accessibility albeit with some clone-specific changes (Figure 3B). Thus, while each knockout line arose from an independent CRISPR ablation, most changes in chromatin accessibility are shared and concordant between the two H1.4-deficient lines, suggesting a potential role for H1.4 in both maintaining open chromatin as well as preventing the opening of chromatin.

**Figure 3.**
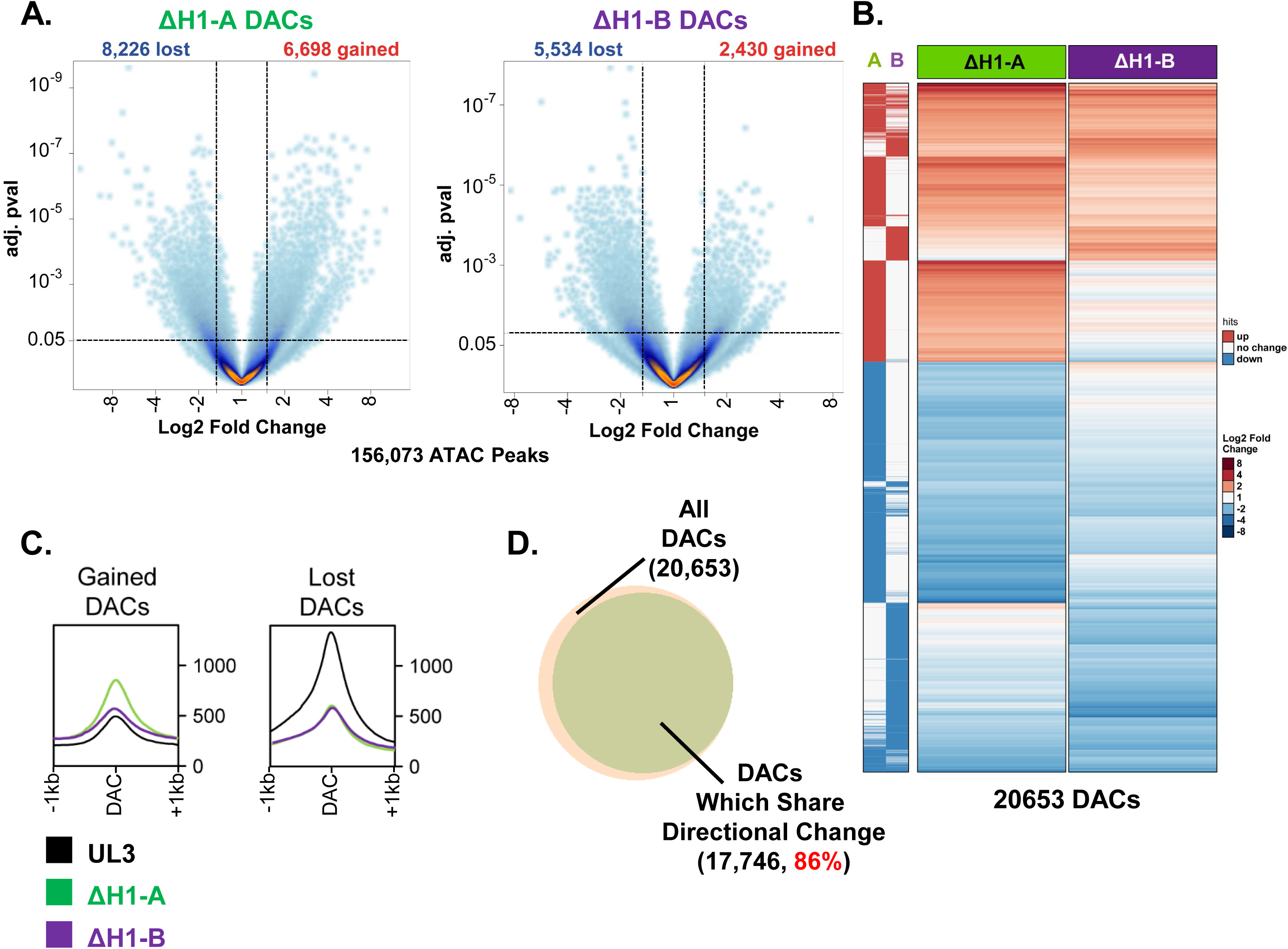
H1.4 ablation results in widespread chromatin reorganization. **A.** ATAC-seq data from two ΔH1.4 clones (ΔH1-A and ΔH1-B) represented as volcano plots. Dashed lines indicate threshold parameters for calling peaks (adj p-val <0.05, fold change >1.5) as differentially accessible. The number of DAC peaks called as opening (red) or closing (blue) is listed above each plot. The total number of peaks represented indicated at the bottom. **B.** Heatmap displaying 20,653 DAC peaks called between ΔH1.4 clones ΔH1-A and ΔH1-B relative to parental UL3 cells. Binary directional changes are summarized (left columns) adjacent to accessibility fold change (right columns) for each DAC peak. **C.** Metaprofile graphs of accessibility data +/-1kb around the 20,653 DAC peaks called as differentially accessible in the ΔH1-A and ΔH1-B clones, separated by direction of change (8,336 gained, left; 12,285 lost, right). Metaprofiles are centered on the peak center as depth normalized reads, and shaded regions above and below each line indicate one standard error to the mean value. Accessibility for the UL3 parental cells as well as each ΔH1.4 clone are displayed (32 discordant DACs are not shown). **D.** Venn diagram illustrating the agreement between the total number of DACs (n=20,653) called in any cell line (peach) and the number of DACs which share the same direction of change in both ΔH1.4 clones (n=17,746, 86%). DAC peak numbers and percentages are indicated.

Having identified a high-confidence set of H1.4-dependent DACs (n=20,653), we examined the metaprofile of the ATAC-seq data at the DACs. First, both gained and lost DACs are small, usually less than 1kb. The accessibility changes at the center of the DAC, but as one moves even 500bp from that center the accessibility goes down and is nearly at baseline 1kb from the DAC center (Figure 3C). We did not observe large domain-level changes in accessibility and instead identified chromatin changes typical of enhancers and other regulatory sites at which transcription factors could mediate the changes. The height and focus of ATAC-seq around the in the UL3 parental peaks are reminiscent of nucleosome positioning at TSSs where an open region is flanked by well positioned +1/-1 nucleosomes. Next, we investigated the differences in regions which gained or lost accessibility. Sites which gain accessibility appear mostly chromatinized in the UL3 parental cells and gain a moderate degree of ATAC-seq signal at the center of the DAC. The gained DACs also had more heterogeneity, with more identified in the ΔH1-A line and this line showing more ATAC-seq signal at the DAC than the ΔH1-B line consistent with more gained DACs in the ΔH1-A line and this may explain some of the cell-line specific phenotypes observed. This suggests that regions with greater DAC peak accessibility experience a mild passive relaxation of chromatin structure with the loss of H1.4, as opposed to an active removal of nucleosomes. The metaprofile of UL3 cells at DACs which lose accessibility shows sites with a high degree of open chromatin, and both cell lines phenocopy the degree of change in ATAC-seq at these lost DACs (Figure 3C). The strong and consistent closure of chromatin at DACs with lower ATAC-seq signal suggests that loss of H1.4 at these sites induces a more active chromatin closure process consistent with H1.4 stabilizing the chromatosome structure, The 32 discordant DACs can be viewed in Supplemental Figure 5D.

### Additional epigenetic changes are concomitant with changes in chromatin accessibility

We again turned to meta-data analysis to further examine the differences between the two ΔH1.4 clones, using subtraction to remove the UL3 cell signal (Figure 4A top panels) and created meta-data plots and difference heatmaps (Figure 4A bottom panels). As ΔH1-A had more DAC peaks, difference heatmaps of the ATAC-seq data were sorted vertically by the amount of change in the ΔH1-A clone. White indicates no change in ATAC-seq relative to the parental, and as such, the DAC peaks show considerable change in accessibility with the loss of H1.4. Loss of accessibility at DAC peaks and shape of these DAC peaks is broadly consistent between the two ΔH1.4 clones. Gained DAC peaks appear more clone specific, with less change apparent in the ΔH1-B line. As seen on other analysis, the ΔH1-B ATAC-seq data are consistent with that from the ΔH1-A line, albeit the fold change in accessibility is somewhat diminished. Raw coverage heatmaps, as opposed to the difference plots, are shown in Supplemental Figure 4. We conclude that the loss of accessibility is widely shared in the two H1.4 knockout lines while the sites where accessibility is gained may be more cell-line specific albeit reproducible within a cell line.

**Figure 4.**
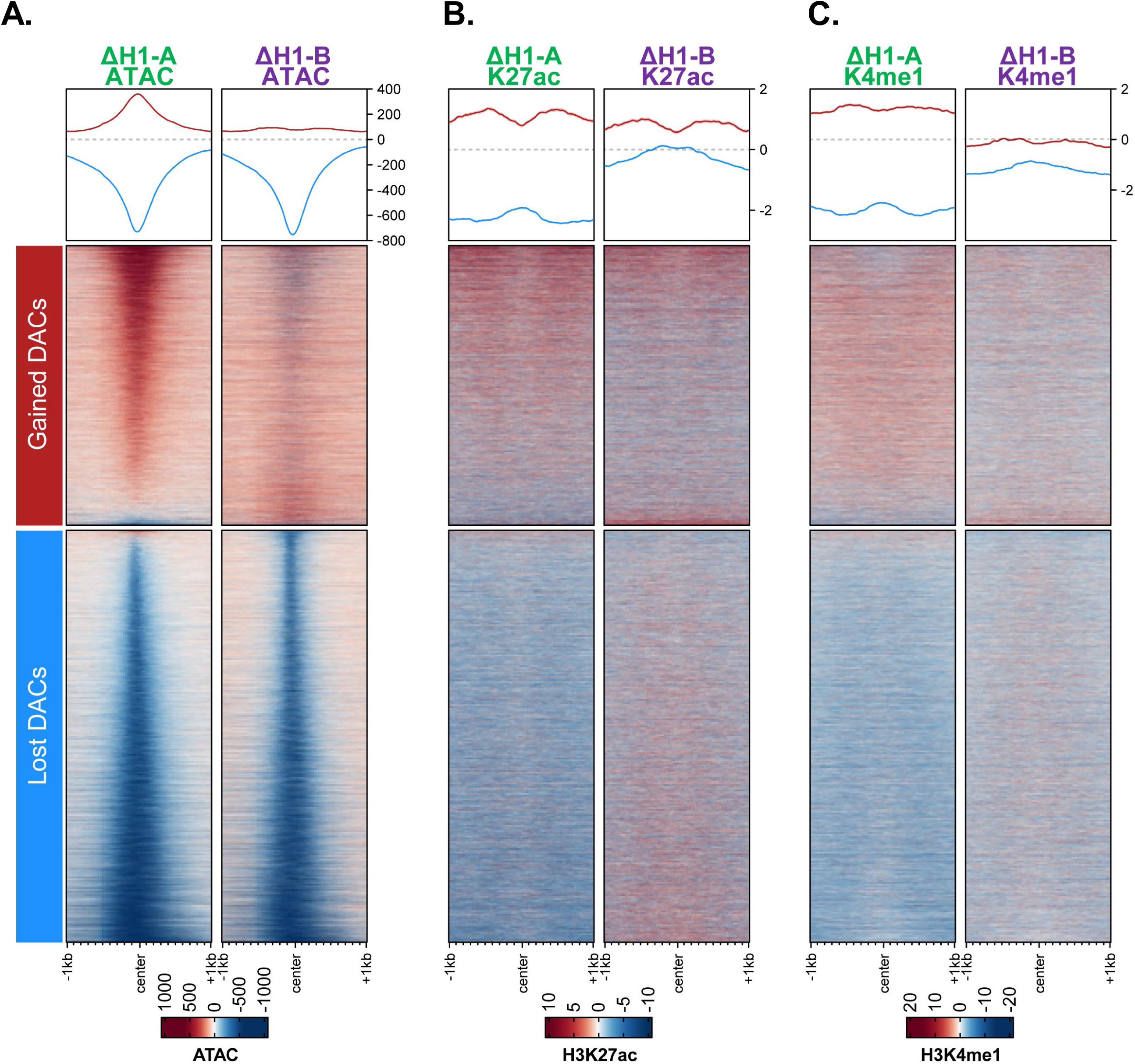
H1-dependent changes in genomic accessibility are accompanied by changes in active histone marks. Difference heatmaps displaying signal (**A**, ATAC-seq; **B** H3K27ac ChIP-seq; **C**, H3K4me1 ChIP-seq) in each ΔH1.4 clone for the combined 20,653 DAC peaks called between the ΔH1-A and ΔH1-B clones. Each row represents a single DAC peak, rank ordered by degree of DAC change in clone ΔH1-A. Signal from the UL3 parental dataset has been subtracted. Red signal indicates gained signal (more open chromatin, 8,336), whereas blue signal indicates less ATAC-seq signal (more closed chromatin, 12,285). Metaprofiles are graphed above each heatmap for data separated by direction of accessibility change with a shaded region above and below each metaprofile to indicate one standard error.

We next examined epigenetic changes occurring on the core nucleosome because of H1.4 loss. ChIP-seq for H3K27ac, a chromatin mark typically found at sites of active transcription (69,70), correlated well to our 20,653 DAC peaks using differential heatmaps. Both ΔH1.4 clones agreed, displaying an increase in H3K27ac at the gained DAC peaks (red line, Figure 4B), and a loss of H3K27ac at the DAC peaks that become more closed (blue line, Figure 4B). These changes in H3K27ac deposition are indicative of a concordant change in chromatin activity and suggest that changes in transcriptional activity might also be occurring in these regions.

Using ChIP-seq, we also examined H3K4me1 deposition in our UL3 parental cells and ΔH1.4 clones to see if loss of H1.4 affected the epigenetic mark typical at cis-regulatory enhancers (69,71). DAC peaks called in the ΔH1.4 clones exhibit roughly equivalent H3K4me1 in both gained and lost DAC peak regions of UL3 parental cells (Supplemental Figure 4C). These same regions display diverging H3K4me1 signal, with regions of gained accessibility gaining H3K4me1, and the opposite for lower accessibility regions of the ΔH1.4 clones (Figure 4C). Together, these findings support our identification of differentially accessible chromatin and agree in the direction of change, whereby sites gaining accessibility also gain active histone marks while sites losing accessibility show a reduction in those same active marks.

### DACs are enriched at enhancers and transcribed genes as well as hetero/quiescent chromatin

The 20,621 DAC peaks were annotated to genomic features and found to intersect primarily extragenic space, with some enrichment observed in introns of annotated genes (Supplemental Figure 5). If anything, there is a de-enrichment of both gained and lost DACs at transcription start sites, suggesting that the regulatory mechanism of these chromatin changes may be in outside gene promoters and may lie in cis regulatory space.

The DACs were then intersected with the Epilogos chromatin state data (https://epilogos.altius.org) to further parse the data. Epilogos annotates the chromatin state in 200bp increments by ChromHMM ChIP-seq data obtained from 127 tissue and cell samples. Each 200bp region is assigned 1 of 6 chromatin states based on the most observed chromatin state in the datasets queried. As in the annotation-based analysis, despite ATAC-seq preferentially capturing transcription start sites, both gained and lost DACs were de-enriched at chromatin typical of promoters (Figure 5A). Despite a dearth of enhancer-like chromatin in the genome (5.8%), we observed a strong enrichment of DAC which lose accessibility with enhancer status (Figure 5A). The sensitivity of enhancers to the loss of H1.4 may be related to the combinatorial mechanisms by which many enhancers can work to regulate promoter activity and transcriptional output. Transcriptionally active chromatin was found to be enriched in gained DACs and depleted of lost DACs, suggesting that transcription may play a role in the opening of the chromatin in absence of H1.4. Again, as observed with the annotation-based analysis, both lost and gained DACs were de-enriched at sites with active or poised TSS chromatin despite the high enrichment of TSSs in ATAC-seq. Finally, and as expected, regions rich in heterochromatin and quiescent chromatin marks were enriched in both gained and lost DACs, suggesting these regions distal from genic space are dependent upon H1.4 for their appropriate compaction, with some regions opening and some closing (Figure 5A). Heterochromatin regions were found to have the greatest gain in accessible chromatin, with a >2-fold increase in the percent of DACs found there relative to the total ATAC peaks. More complete intersection of all 15 chromatin states and four genic states are in Supplemental Figure 5, including DACs which are opposing.

**Figure 5.**
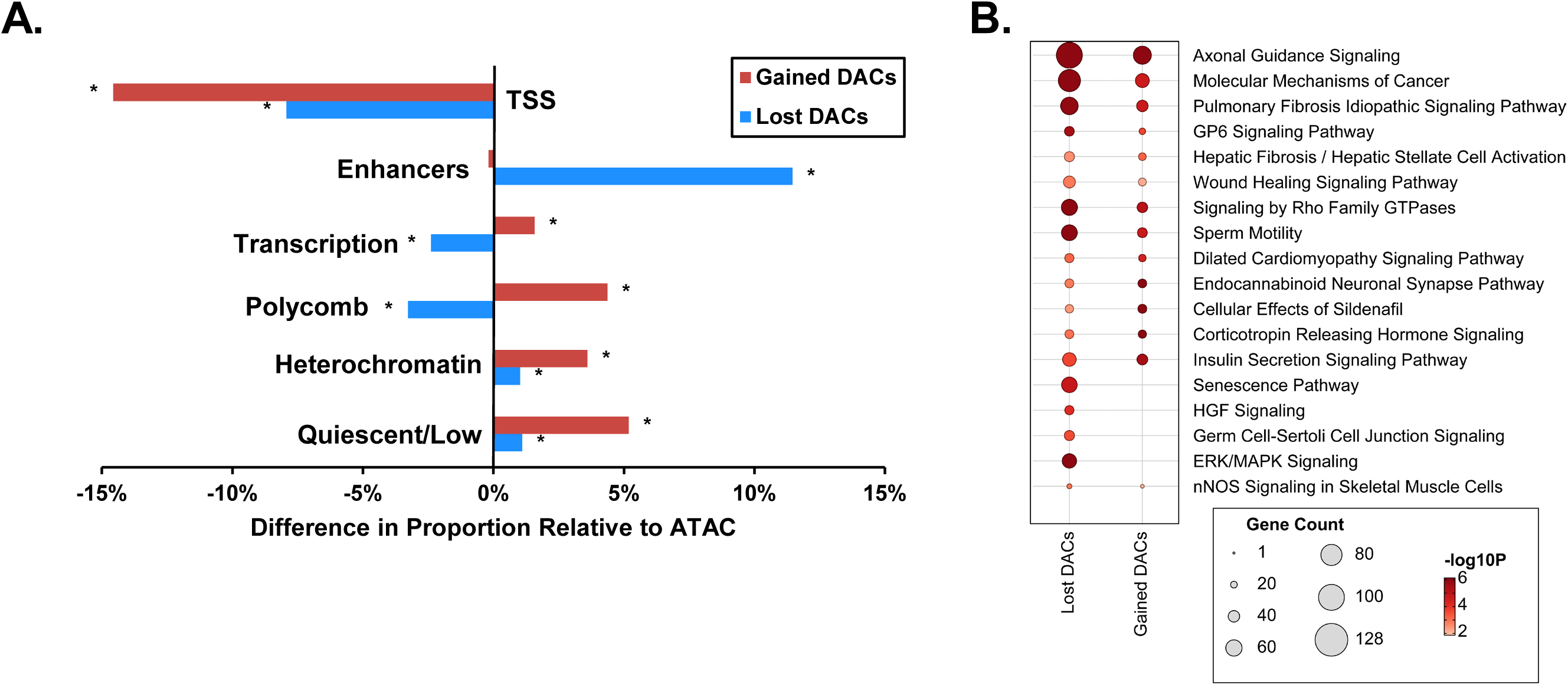
H1.4-dependent chromatin intersect epigenetically regulated regions including transcriptionally active and compacted chromatin. **A.** Proportional difference in proportion of gained DACs (red) and lost DACs (blue) relative to the distribution of all ATAC-seq peaks identified. Each row indicates the enrichment (positive) or de-enrichment (negative) of each of the six Epilogos chromatin states (transcription start site (TSS), enhancer, transcription, polycomb, heterochromatin and quiescent/low) for each genomic 200bp genomic bin when compared to all possible open chromatin, Asterisks indicate bins which meet statistical significance defined by a Pearson chi-squared residual > 2. Pearson residuals 2 and 4 represent approximately ɑ = 0.05, and ɑ = 0.0001 and calculated Pearson residuals are in Supplemental Table 2. Distribution of ATAC-seq and raw counts of gained and lost DACs relative to all Epilogos bins are shown in Supplemental Figure 5. **B.** The enrichment of gene signaling pathways the gained and lost DACs intersect represented as a dot plot. Larger circles indicate pathways with more genes, darker circles indicate greater number of DACs being identified as intersecting genes in the pathway.

To investigate if certain genetic pathways were particularly enriched in H1-dependent chromatin, we performed Ingenuity Pathway Analysis on both the gained and lost DACs by assigning each DAC to a gene, either the gene it intersects or the nearest gene body. We found that the differentially accessible chromatin sites localized to vital signaling pathways related to the regulation of cancer, cell-cell communication and GTPase signaling (Figure 5B). The lost DACs showed a unique enrichment in critical phosphorylation signaling pathways such as ERK/MAPK and HGF/c-MET suggesting these signaling pathways may be unusually dependent upon H1 for proper chromatin organization and activity. These pathways are central to the regulation of responses to stimuli and cell identify in osteocytes (72–74).

### Chromatin regions relying on H1.4 for accessibility are enriched for AP-1 transcription factors

Observing the enrichment for intronic and extragenic enhancers and other cis-regulatory chromatin states in ΔH1.4 DAC peaks, we asked which transcription factor motifs are located at the DACs to determine if any TF might mediate these changes (MEME suite). We first interrogated transcription factor motifs in the DACs with lower accessibility. Strikingly, we observed a strong enrichment in motifs for the various members of the Activator Protein 1 (AP-1) transcription factor family in both ΔH1.4 clones (Figure 6A). AP-1 is a heterodimer of Fos and Jun-like transcription factors, and often serves as an intracellular intermediate in critical signaling cascades (75–77). We were curious to know if AP-1 transcription factors have been previously reported to bind to sites we identified as closed ΔH1.4 DAC. Using the ENCODE ChIP-seq data for AP-1 transcription factors, we plotted AP-1 ChIP-seq signal around the DAC peaks (78). The ChIP-seq signal for Jun, JunD, Fos and FosL1 correlates well with the motif analysis, exhibiting as a high and focused peak of AP-1 transcription factor binding at the center of the closed ΔH1.4 DACs.

**Figure 6.**
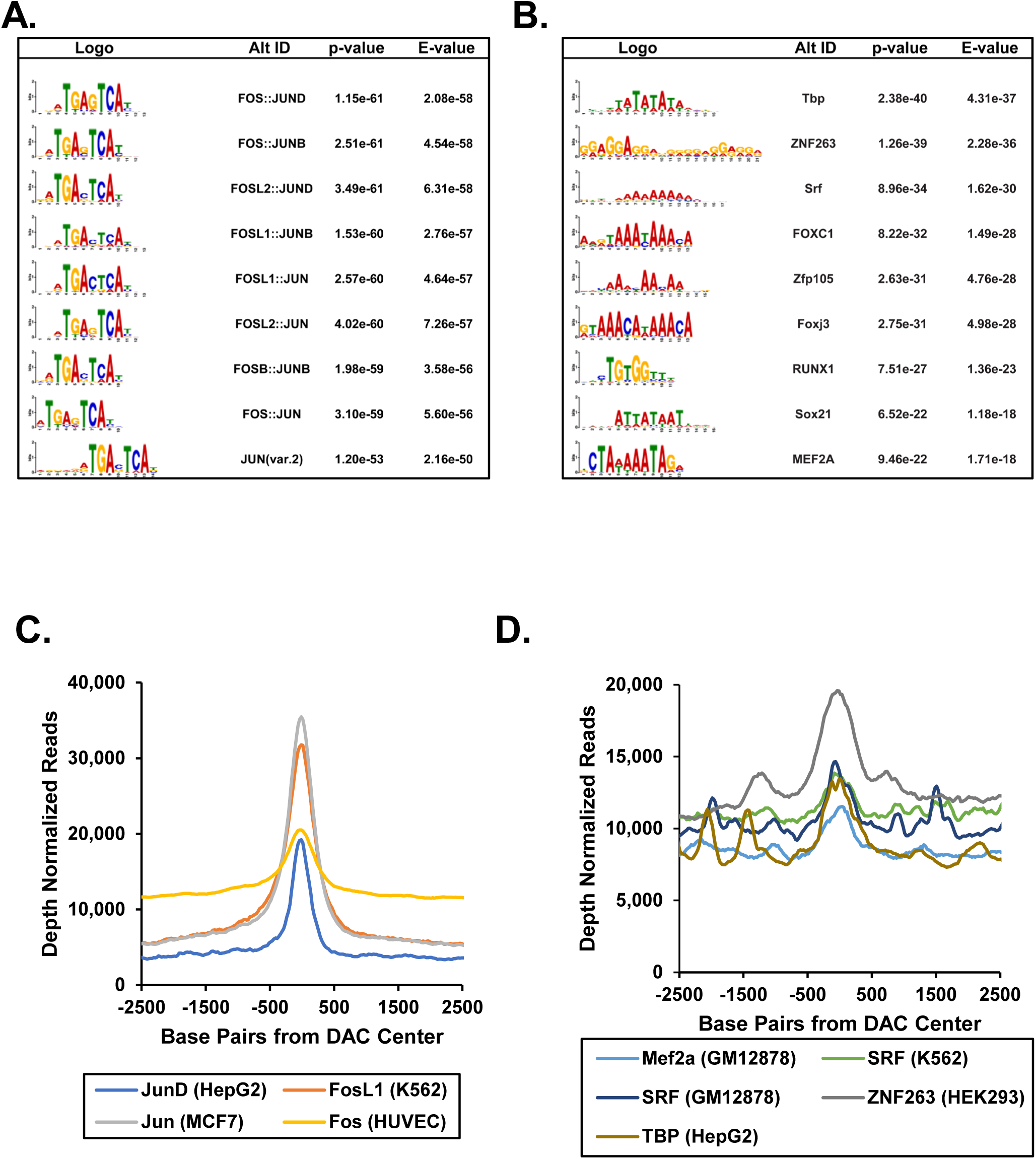
Motif Analysis suggests gained DACs may be interacting with basal transcription machinery while reduction in AP-1 family may be responsible for the lost DACs. Transcription factor motif enrichment and in vivo binding were analyzed for ΔH1.4 DAC peaks. MEME analysis was performed on the 12,285 DAC peaks with lower accessibility (**A**) and the 8,336 DAC peaks which gained accessibility (**B**) and enriched LOGOs are shown. The ID of the motif and both a p-value and E-value representing the relative enrichment of each motif are shown. Metaprofiles of the available ENCODE ChIP-seq data for transcription factors with enriched motifs were plotted +/-2.5kb around the center of each DAC peak for those motifs identified at the lost DACs (**C**) and the gained DACs (**D**)

Although we did not observe enrichment of a specific transcription factor family with the gained DAC peaks, we did observe an enrichment of motifs for broadly expressed chromatin-sampling transcription factors (Figure 6B), such as TBP and SRF. These observations suggest a potential mechanism where nucleosome structure weakened by loss of H1.4 could be further opened by sampling transcription factors and possibly even RNA polymerase, as predicted by the increases in H3K27ac at these DACs. To determine if these commonly expressed transcription factors could bind these gained DACs, we plotted ChIP-seq data from ENCODE (78) around our gained DACs and observed a small but significant peak of binding at the center of our identified locations for Mef2a, Srf, Znf263 and TBP, all factors which have binding motifs within the gained DACs (Figure 6D). Similar binding of these basal factors was not observed at 30,000 randomly selected genomic locations (Supplemental Figure 6A) nor was biding of the AP-1 observed at these random sites (Supplemental Figure 6B).

### H1.4 loss results in profound dysregulation of nascent transcription

As we had selected for stable ΔH1.4 cell lines, it is possible that loss of H1.4 could result in compensatory mechanisms that limit our ability to detect DEGs by standard RNA-seq (e.g., changes in RNA stability). Metabolic labelling of RNA with 4-thio-uracil (4sU) allows for the measurement of transcriptional changes distinguished from changes in RNA maturation and stability. Live UL3 and ΔH1.4 clones were exposed to 4sU for 10 minutes, briefly labeling all newly transcribed RNAs, which were subsequently harvested and used to generate sequencing libraries for 4sU-seq (56–59, 83). While only 6-12% of genes change when measured by RNA-seq, 4sU-seq reveals that loss of H1.4 has a much broader effect on raw transcription such that 9-38% of expressed genes (ΔH1-A 1,357 DEGs, ΔH1-B 6,003 DEGs). In both ΔH1.4 cell lines, there was a bias towards downregulation of DEGs (Figure 7A), and 56% of the DEGs were shared between the two knockout lines (Figure 7B). As observed with RNA-seq, 4sU-seq DEGs are in directional agreement (Figure 7C). We further validated the 4sU-seq by performing RT-qPCR on a small subset of the DEGs including the ΔH1-A specific DEG INHBE (Supplemental Figure 7). We used the more sensitive 4sU-seq results to examine expression changes in the transcription factors predicted to bind the DACs. Expression of the AP-1 transcription factors were repressed in both ΔH1.4 clones, however each cell line exhibited a different AP-1 family member dysregulation or had closely related genes similarly dysregulated (e.g., FosL1 and FosL2). (Figure 7E). We did not observe significant expression changes in the transcription factors predicted to bind the gained DACs, suggesting that if these factors were mediating the changes, the mechanism may be independent of their expression levels and instead may involve their DNA sampling activities (Figure 7D).

**Figure 7.**
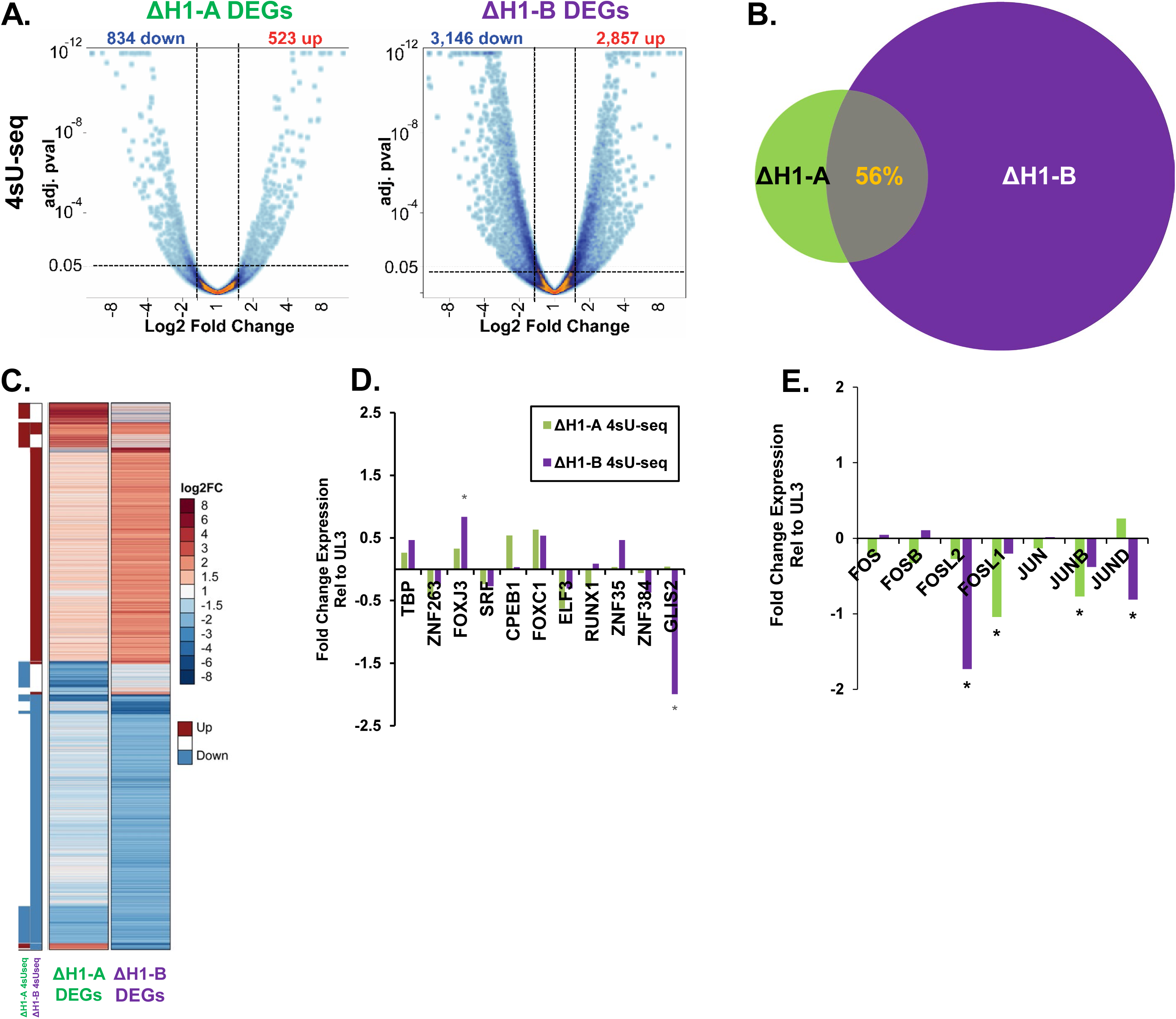
H1.4 ablation results in robust changes in nascent gene transcription. **A.** 4sU-seq data from two ΔH1.4 clones (ΔH1-A and ΔH1-B) represented as volcano plots. Dashed lines indicate threshold parameters for calling peaks (adj p-val <0.05, fold change >1.5) as differentially expressed. The number of upregulated genes indicated in red and downregulated in blue are listed above each plot. **B.** Venn diagram illustrating the agreement in DEGs identified in the ΔH1-A (green) and ΔH1-B (purple) ΔH1.4 CRISPR/Cas9 clones. **C.** Heatmap displaying the 4sU-seq coverage data for the 7,103 genes called as differentially expressed in the 4sU-seq experiments between ΔH1.4 clones ΔH1-A and ΔH1-B relative to parental UL3 cells. Binary directional changes are summarized (left columns) adjacent to accessibility fold change (right columns) for each gene. Changes in expression of transcription factors in the ΔH1-A (green) and ΔH1-B (purple) ΔH1.4 clones found to bind the gained DACs (**D**) and lost DACs (**E**) are shown using the 4sU-seq data are shown relative to UL3 expression. Asterisks indicate genes which reach the statistical significance thresholds set.

Although both ΔH1.4 clones have high similarity between DEGs, the directionality of DEG change is not always preserved between clones. We next investigated our 4sU-seq DEGs for common pathways (IPA, Qiagen) in hopes to better understand the phenotype of the ΔH1.4 clones. Surprisingly, and despite differences in DEG directionality, both ΔH1.4 DEG sets were enriched for prototypic pathways for osteosarcoma cells, such as cell proliferation, DNA damage, receptor activator of nuclear factor-κB ligand (RANKL) signaling, and integrin signaling as similarly dysregulated (Figure 8A). These pathways are central to osteocyte cell signaling and more importantly contain pioneer factors that could interact with H1 linker histones (72–74, 79). Further examination of pathway gene set overlap reveals that a specific gene family, the Immediate Early Genes (IEG), is driving the dysregulation in processes impacted by H1.4 loss (Figure 8B). IEGs are the primary mediators of extracellular signaling pathways, often involved in stress or immune responses (80–82). IEGs are highly regulated during early transcription elongation by transcriptional pausing, a mechanism that allows for rapid and robust gene activation by pre-loaded RNA Pol II at the promoters (83–85). A role for H1 regulating immediate early genes has been described as working through an interplay between PARP-1 and H1 linker histones (86). This suggests the intriguing possibility that H1.4 is plays a regulator role during early transcriptional elongation, possibly during transcriptional pausing.

**Figure 8.**
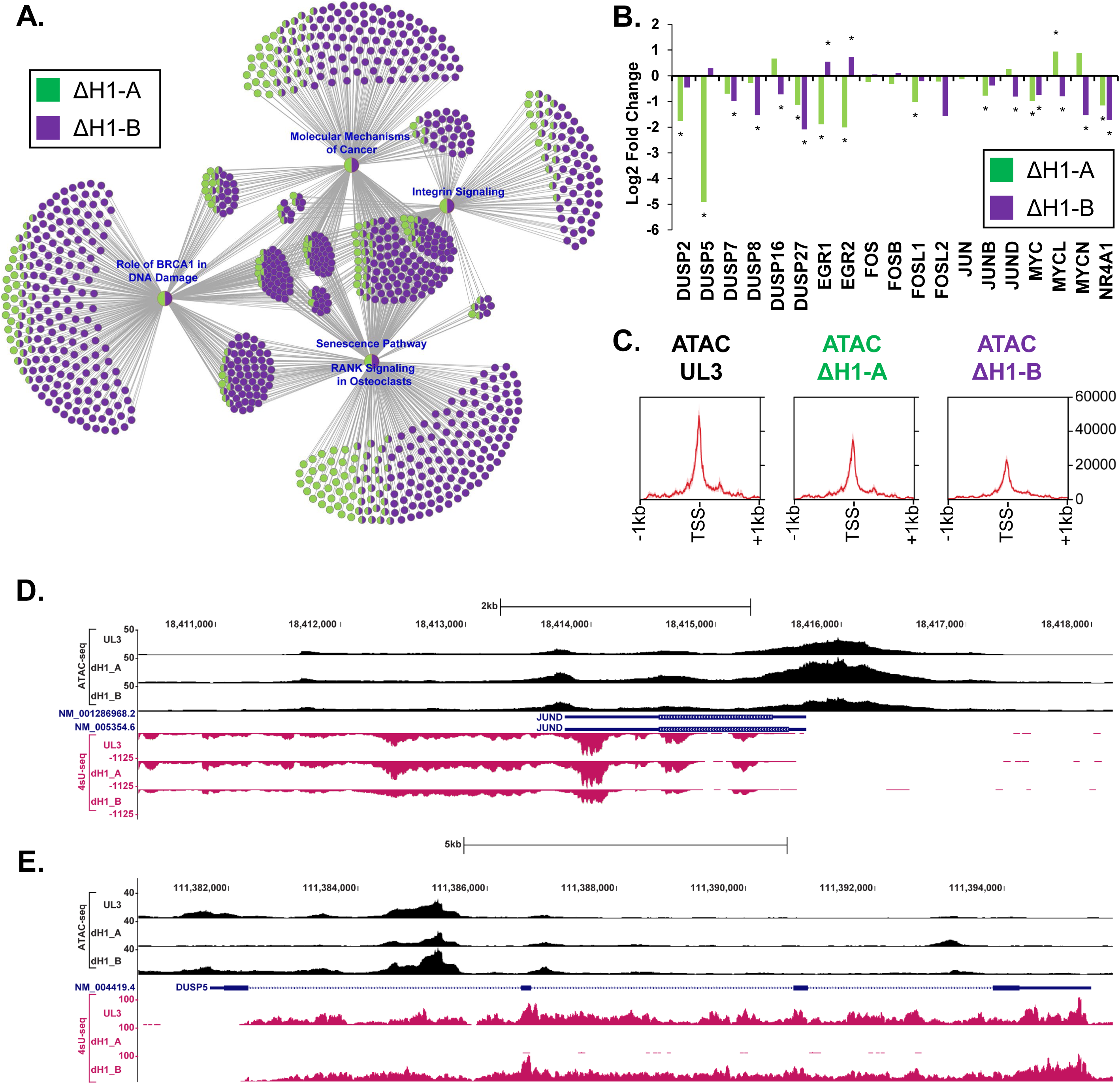
H1.4-KO cell lines show dysregulation of similar pathways related to cell growth and responsiveness. **A.** Ingenuity Pathway Analysis of the 4sU-seq DEGs from ΔH1-A (green) and ΔH1-B (purple) ΔH1.4 CRISPR/Cas9 clones visualized as a Category Network (CNET) plot, with shared DEGs showing both colors. Each circle represents a gene and nodes of signaling are indicated with their name in blue. Dysregulated genes shared across several pathways are indicated by multiple grey lines connecting them to various signaling nodes. **B.** Changes in expression of the Immediate Early Genes in in the ΔH1-A (green) and ΔH1-B (purple) ΔH1.4 clones are shown using the 4sU-seq data and are shown relative to UL3 expression. Asterisks indicate genes which reach the statistical significance thresholds set. C. Metagene of ATAC-seq data from the UL3 parental cells (Black), the ΔH1-A (green) and ΔH1-B (purple) ΔH1.4 CRISPR/Cas9 clones +/-1kb around the 61 expressed Immediate Early Genes. Metagenes are centered on the TSS of each IEG with the promoter pointing to the right in all cases. D. Coverage of ATAC-seq (black) and 4sU-seq (rose) data at the JunD locus. JunD is on the negative strand and is upregulated in the ΔH1-A clone and shows increased promoter accessibility. E. Coverage of the ATAC-seq (black) and 4sU-seq (rose) data in the Dusp5 locus. Dusp5 is encoded on the positive strand and expression is lost in the ΔH1-A clone and shows decreased promoter accessibility.

Finally, we examined the 61 IEG promoters previously identified for changes in ATAC-seq and observed a marked loss of promoter accessibility in the ΔH1.4 clones relative to parental UL3 cells (Figure 8C). Examination of ATAC-seq coverage data for the JunD and Dusp5 promotors is consistent with this loss of accessibility (Figure 8 D&E). In these gene coverage maps, the 4sU-seq data is also displayed, indicating both the un-spliced nature of these data as well as their sensitivity for changes in transcription. Based on these data, it appears the broad family of critical Immediate Early Genes is sensitive to perturbation of H1.4 levels in these cells and suggests H1.4 may play a role in keeping these promoters open during early RNA Pol II elongation.

## DISCUSSION

Historically, the role of linker H1 has focused on its role in the compaction of the chromatin fiber within the nucleus, and it was mostly thought of as a structural protein (11). However, work from our lab and others has shown the role histone H1 plays in the regulation and reorganization of the mouse mammary tumor virus promoter, specifically in response to glucocorticoids (21–25, 87–102) as well as the withdrawal of corticosteroids from cells. Moreover, histone H1 has been shown to dominantly repress the totipotent state of embryonic stem cells (103). This previous work highlights the specific structural changes, notably phosphorylation and eviction, that H1 can play in the direct regulation of gene activity and signaling responses.

In this work, we established five genetically distinct H1.4-deficient cell lines and observed very similar compensatory mechanisms (e.g., upregulation H1.1, H1.2, H1.0) as well as the dysregulation of shared response pathways such as TNFα/NF-κB and integrin/ECM signaling. We then identified over twenty thousand individual sites with changed chromatin architecture termed DACs. These DACs were usually less than a kilobase in length, which lends them more to localized changes in chromatin activity rather than large-scale genomic rearrangements based on chromosomal compaction. These changes in chromatin architecture are concomitant with changes in nucleosomal histone modification, with regions losing accessibility also losing H3K4me1 and H3K27ac marks, and the reverse being true for sites with gained accessibility also gaining epigenetic marks consistent with gained activity. Despite ATAC-seq’s bias towards capturing the most open regions of the genome such as promoters, we observed a relative dearth of promoter-proximal DACs and instead observed most DACs are located extragenically. Many DACs were observed in the heterochromatin and quiescent chromatin regions, consistent with the compaction role of H1 in genome stability. Regions with transcriptionally active chromatin were found to be enriched in gained DACs and de-enriched in lost DACs, suggesting a potential interplay between transcriptional activity and chromatin accessibility in the absence of H1.4. Lost DACs were found to have a strong tropism for enhancer regions, both intronic and extragenic. Indeed, the identification of enriched cis regulatory elements and transcription factor binding sites suggests a novel role for H1.4 in gene regulation that deserves further exploration (Figure 9).

**Figure 9.**
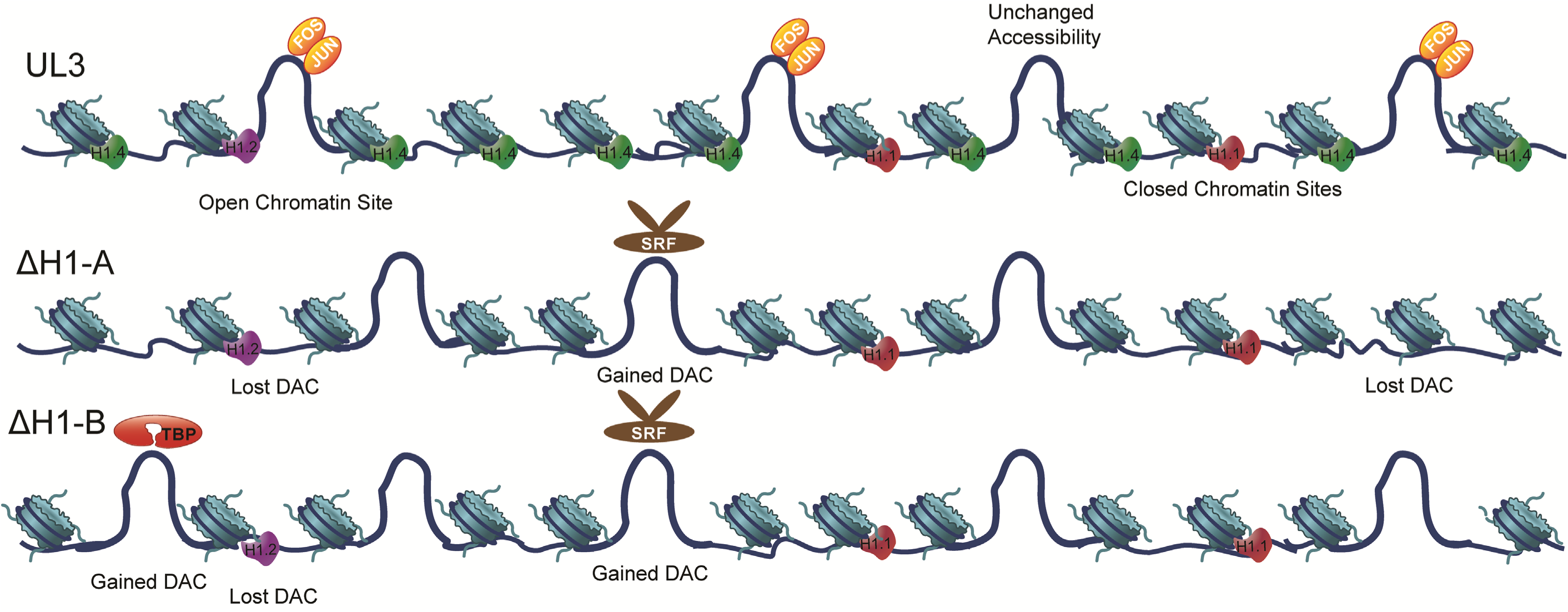
Model of Changes to DACs. Changes in chromatin accessibility are observed when a cell line which expressed high levels of H1.4 has this H1 variant ablated. Genetically distinct cell lines show remarkably similar changes in accessibility with this genetic ablation, especially at sites which lose accessibility. Here, the UL3 parental line expresses a mixture of H1 variants which includes H1.1 (red), H1.2 (purple) and H1.4 (green). The globular domain of H1 linker binds the histone octamer at the nucleosomal dyad and contacts the DNA located here as well as the 10bp extensions of linker DNA either entering or exiting the nucleosome to form the chromatosome. At many of the sites observed to lose accessibility in the ΔH1-A and ΔH1-B lines contain binding motifs for the transcription factors Fos and Jun (AP-1 family members) and AP-1 family members have been shown to bind these sites in vivo. The knockout cells also show reduced expression of AP-1 family members, albeit in a cell-line specific manner (e.g., Fos and JunB in ΔH1-A but FosL1 and JunD in the ΔH1-B line). The H1.4-deficient cells also exhibit gain of accessibility phenotypes. These gained sites coincide with motifs for more commonly expressed transcription factors such as TBP (red) and SRF (brown) which, in the absence of H1, may sample the DNA more and facilitate the opening of chromatin. In some cases, the effects of the loss of H1.4 may be alleviated by increases in H1.1 or H1.2 at sites hitherto occluded by H1.4.

The loss of H1.4 had a signification impact on transcription in both ΔH1.4 clones, with many DEGs being clonal common, and sharing similar cell signaling pathways. We identified IEGs, a set of genes regulated by transcriptional pausing, as highly impacted by the loss of H1.4 and observed a loss of accessibility at these promoters. Although DEG sets in both ΔH1.4 show significant overlap, the direction of DEG change was dissimilar, especially the IEGs. For example, Egr1, Egr2 and FosL1 showed strong downregulation only the ΔH1-A line while FosL2, MycN and NR4A1 showed reduced expression in the ΔH1-B line (Figure 8B). We propose a role for H1.4 in gene regulation, where H1.4 is necessary to maintain appropriate regulation of early transcription elongation, possibly during transcriptional pausing. Alternatively, H1 has been shown to play a repressive role at specific genes, and eviction of H1 is seen to have a positive effect on gene expression. It is possible that loss of H1.4 impacts how pioneer factors, such as FOXA or PARP interact with chromatin, resulting in changes to chromatin accessibility and gene expression. We observed AP-1 motif enrichment at DAC peaks that close upon loss of H1.4. Reports are mixed as to whether AP-1 family members can act as pioneer factors (101–103). However, as AP-1 binding has been shown to be highly collaborative with other transcription factors (104), perhaps cell-specific factors may coordinate with AP-1 to control chromatin accessibility via a transcription factor with pioneer activity (e.g., GATA, RUNX). The identification of AP-1 family members as unusually sensitive to H1.4 levels also raises the possibility that the NF-κB signaling networks may depend upon H1.4 for proper regulation or potentially hinting at a role for H1 interplay with these mitogens. This is consistent with the DEGs observed here by with RNA and 4sU-seq. Additionally, while H1 has been shown to play a structural role in the condensation of chromatin for DNA replication, and thus plays an important role in the cell cycle, here we observed differential chromatin to intersect gene loci which participate in ERK/MAPK and HGF/c-MET signaling, pointing for a more direct role of H1 in the control of the expression of these genes or the potentiation of these signaling networks.

Loss of regulatory control of enhancers, transcriptional pausing or chromatin accessibility could result in the directionally inconsistent changes we observe in gene expression, as well as the specific impacts to the gene expression of highly paused genes (IEGs) that we observe with the absence of H1.4. We can’t rule out the impact of H1 variant compensation in our system, nor can we ignore the clonal differences that could be due to the H1 variant pools differences seen in each ΔH1.4 clone at the protein level. In siRNA experiments, H1.0 has already been shown to play a compensatory role when H1.4 is lost (27). However, the dramatic changes we observe to both chromatin accessibility and gene expression are robust and suggest new transcriptional regulatory roles for H1 that have previously not been reported. More work is needed to determine the mechanistic role H1 plays at immediate early genes, as these promoters are relatively free of H1 under both naïve and stimulated conditions (98). There are potentially H1-depenent enhancers which regulate these genes, which begs the question as to how H1 might regulate enhancer activity and accessibility. It is similarly possible that H1 is important for properly positioning the RNA Pol II machinery prior to activation (88) and in its absence the organization of trans-acting factors breaks down. Alternatively, perhaps H1 plays a role in the regulation of upstream antisense or bidirectional transcription, a common feature of IEGs (106,107). These and additional possibilities represent tractable questions in our genetically modified cell models and the promise of further insights into the biological function(s) of linker histones (104).

## DATA AVAILABILITY

Tandem mass spectrometry data as well as ThermoFisher Proteome Discoverer analysis results are available at PRIDE. Next generation sequencing data can be found at Gene Expression Omnibus GEO. Custom bioinformatic tools can be found on GitHub (https://github.com/jmw86069)

## FUNDING

The work described here was supported by funding from the National Institute of Environmental Health Sciences (Z01-ES071006-23).

## Supporting information

Supplemental_Table_1_PCR_Primers

Supplemental_Table_2_Epilogos_Residuals

## ACKNOWLEDGEMENT

We acknowledge the passing of our friend and colleague Dr. C. David Allis and his significant contributions to the fields of chromatin biology and epigenetics and his prescient insights into the importance and contributions of the linker Histone, H1.

We would like to thank the members of the Archer lab, David Fargo and Adam Burkholder for their help in NGS data analysis, Paul Wade and Jason Watts for their internal review of this manuscript. We would specifically like to thank Ginger Muse for her critical reading and suggestions. We would like to thank Artiom Gruzdev of the NIEHS Gene Editing and Mouse Model Core as well as members of the NIEHS Epigenomics and DNA Sequencing Core Facility for their technical expertise.

## CONFLICT OF INTEREST

The authors declare no conflicts of interest.

---

**Supplemental Figure 1:**
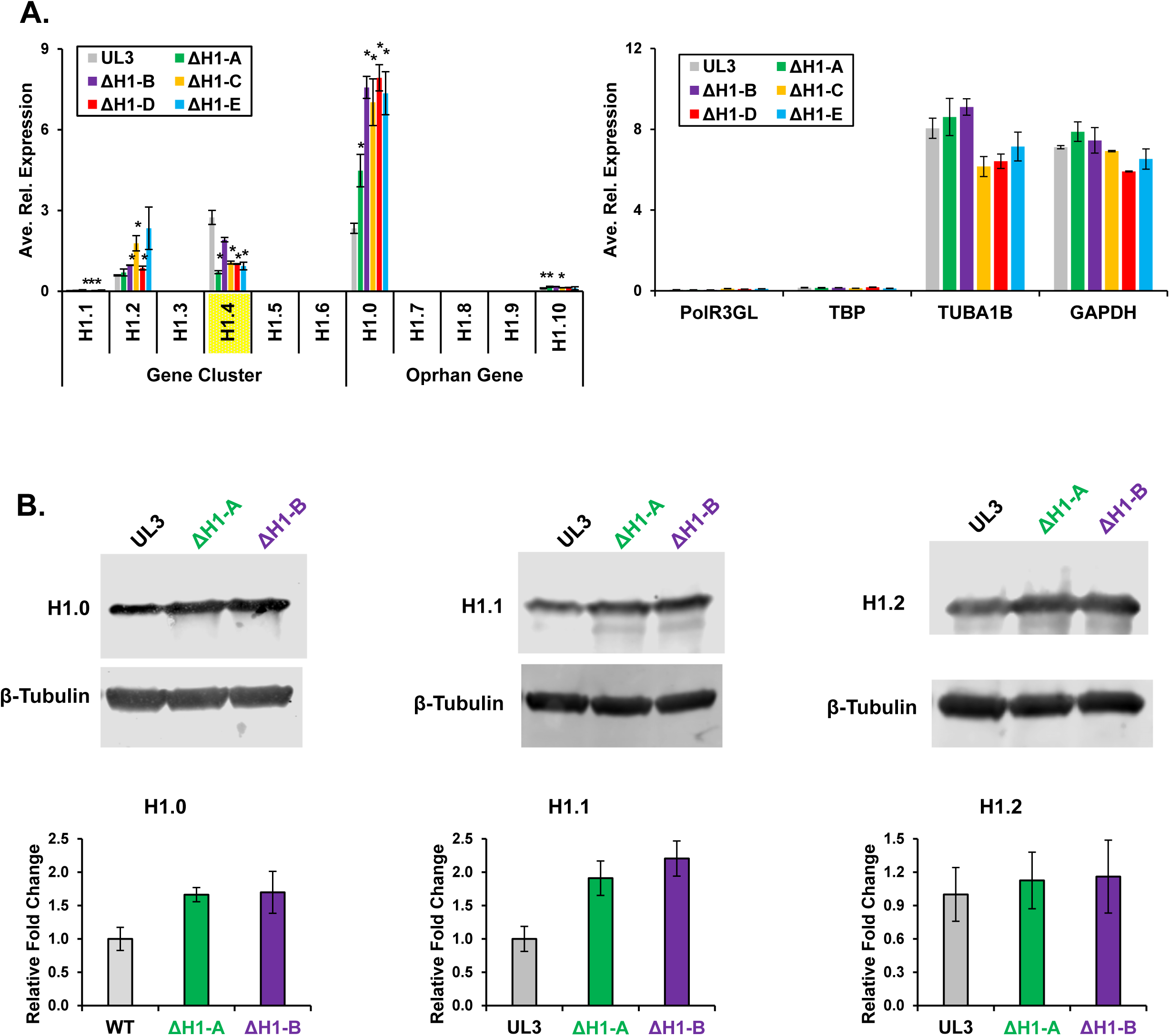
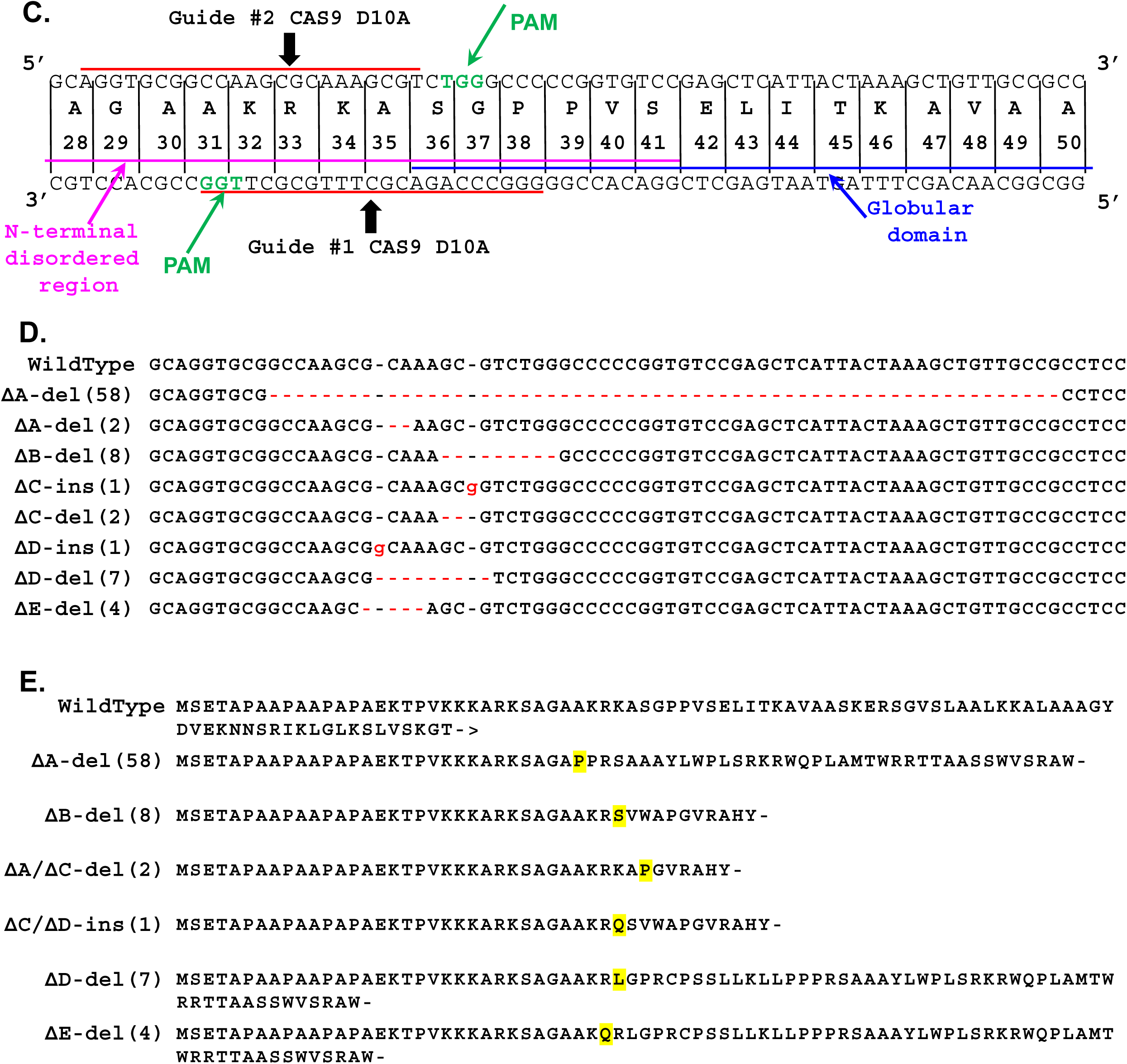
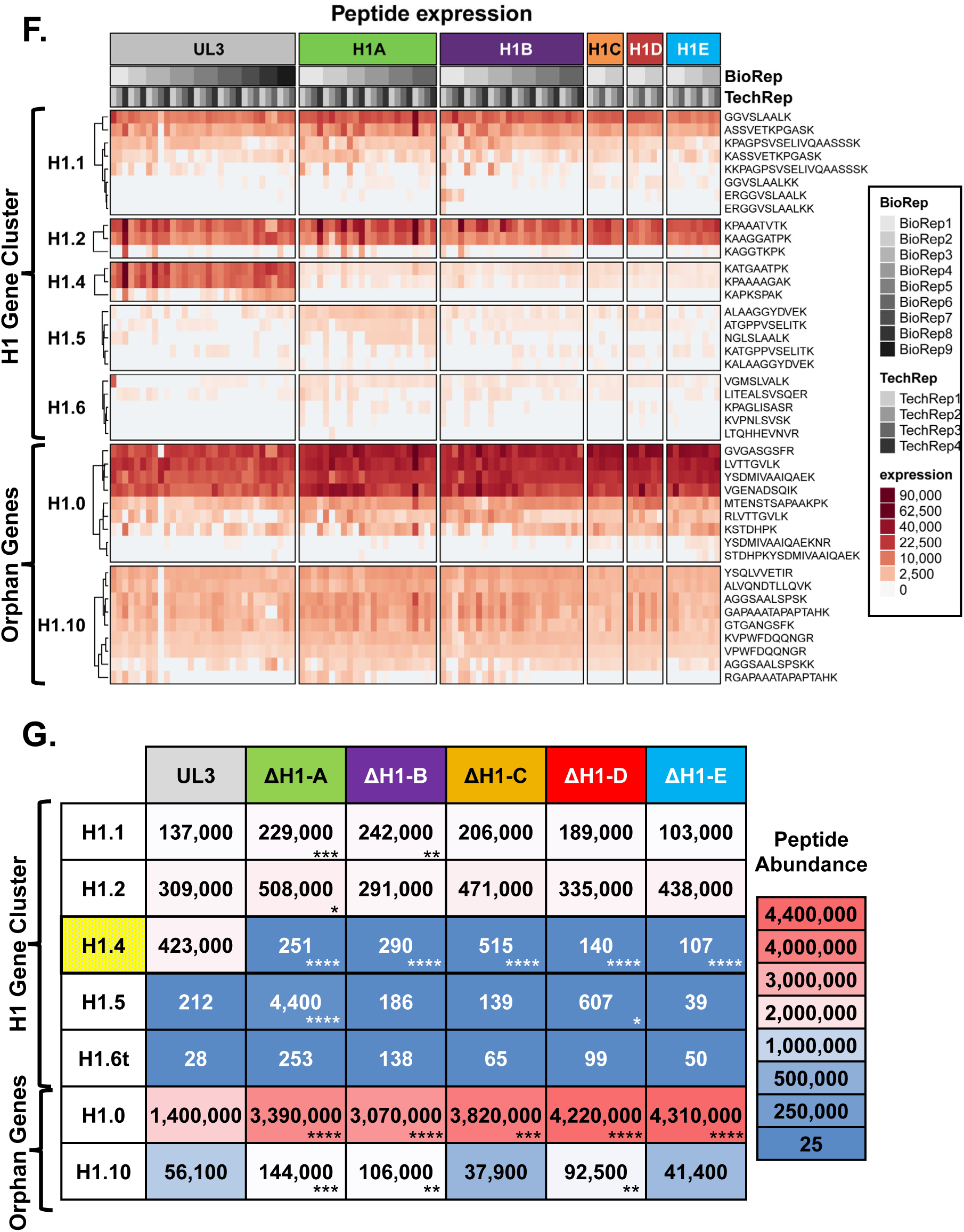
CRISPR ablation of the H1-4 gene and MS/MS of H1 Variants. **A.** RT-qPCR results of relative expression in the parental UL3 cells of H1 gene cluster and orphan gene H1 variants (left) and loading controls genes (right) for parental UL3 cells and five ΔH1.4 CRISPR/Cas9 clones. Error bars indicate standard deviations and asterisks indicate p-values<0.05 in a student’s t-test. **B.** Western blot visualizing changes in the H1.0 (left), H1.1 (center) and H1.2 (right) variants (top panels) relative to β-tubulin loading control (lower panels) in the ΔH1-A and ΔH1-B clones relative to UL3 cells. Densitometry quantification of variant changes relative to UL3 cells from replicate blots shown below. Densitometry values are relative to tubulin and error bars indicate confidence value 95%. **C.** Graphical representation of the CRISPR/Cas9 sgRNA targeting strategy used for the H1-4 gene locus. The relevant H1-4 genomic sense and antisense sequence as well as the translated sequence are depicted with annotations for each sgRNA (red lines), The two gRNA targeting regions of the gene are indicated with red lines, the CRISPR PAM sequences shown in green, the n-terminal disordered region in pink and the globular domain in blue. The two guide locations are labeled in black. **D.** Results of Sanger sequencing of the CRISPR-induced mutations. The wild-type sequence is indicated at the top while the pairs of genomic sequences of the various indel mutations in the knockout lines are indicated below labeled A-E. The ΔH1-B line mutation was homozygous. Red dashes indicate deletion of a nucleoside residue, lower case red text indicate insertions. The number of altered nucleosides is indicated in parentheses with a description of the mutation (del or ins). **E.** Models of the translated amino acid sequences of the parental UL3 and five ΔH1.4 CRISPR/Cas9 clones, labeled A through E. At the top is the wild-type protein sequence, while below are the translated results of the indel mutations. The first altered amino acid is indicated in yellow, with all subsequent amino acids being incorrect, and an early termination codon is indicated with a dash. **F.** Heatmap of relative spectral count data for H1 Gene Cluster (H1.1-H1.6) and Orphan Gene (H1.0, H1.7-H1.10) H1 variants from tandem MS analyses from parental UL3 cells and five ΔH1.4 CRISPR/Cas9 clones. Columns indicate each replicate (bio/technical) and are grouped by cell line; rows represent uniquely observed spectra for each peptide, the amino acid sequence of which is shown to the right. **G.** Heatmap representation of MS/MS spectral count data H1 Gene Cluster (H1.1-H1.6) and Orphan Gene (H1.0, H1.7-H1.10) H1 variants (rows) from analyses from parental UL3 cells and H1.4 knockout lines (columns). Adjusted p-value thresholds for variant changes relative to UL3 cells are indicated with asterisks (*≤0.05, **≤0.01, ***≤0.001 and ****≤0.00001). No unique peptides were observed for variants H1.3, H1.7, H1.8 and H1.9 is an expressed pseudogene in humans.

**Supplemental Figure 3:**
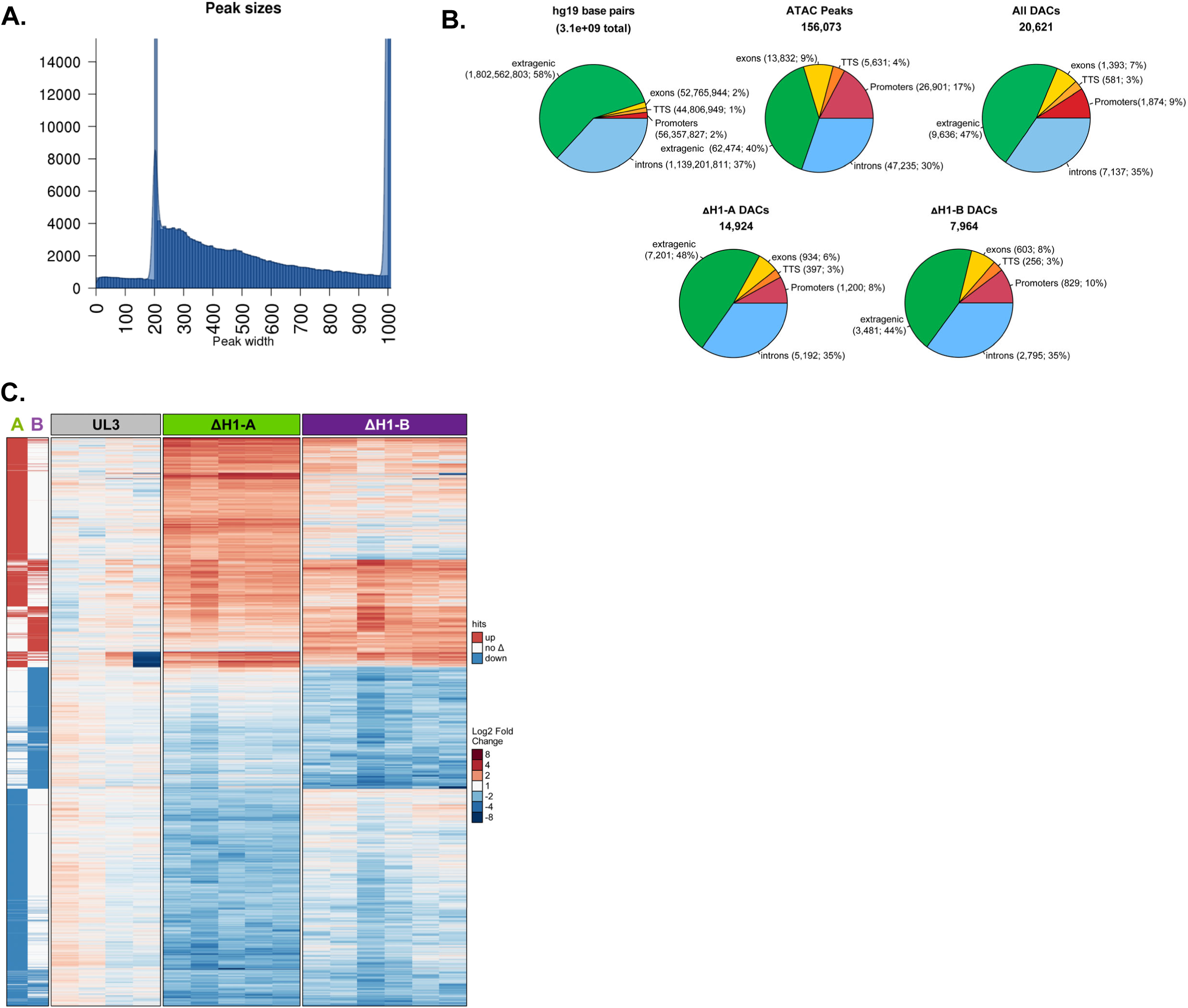
ATAC-seq Peak Analysis and Identification of Differentially Accessible Chromatin Peaks. **A.** Histogram graph of the overall length of all 1kb sliced ATAC-seq peaks called in any cell line. **B.** DAC peak distribution relative to genomic annotations (promoter, exons, transcription termination (TTS), introns and extragenic) in the two ΔH1.4 clones (bottom). Distribution of these features in the genome, amongst all ATAC Peaks and all DACs (top). **C.** Heatmaps displaying ATAC-seq data for each biological replicate at the combined 20,653 DACs called in ΔH1-A and ΔH1-B clones. Binary directional changes are summarized (left columns) adjacent to accessibility fold change (right columns) for each DAC peak.

**Supplemental Figure 4:**
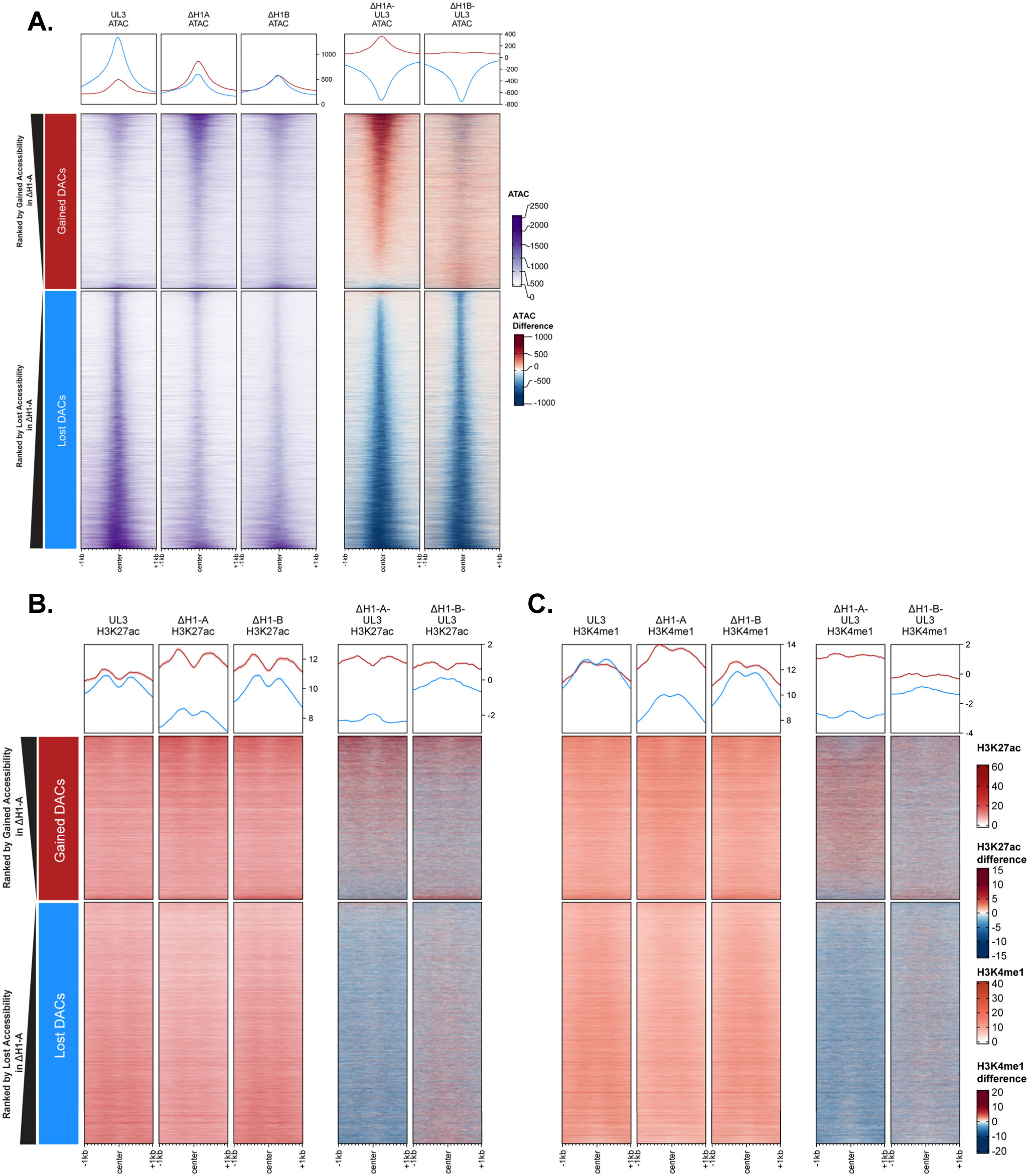
Heatmaps of Chromatin Marks and Epilogos Bins at ATAC-seq DACs. Coverage heatmaps displaying coverage signal (**A**, ATAC-seq; **B** H3K27ac ChIP-seq; **C**, H3K4me1 ChIP-seq) in UL3 parental cells and each ΔH1.4 clone for the combined 20,653 DAC peaks called between the ΔH1-A and ΔH1-B clones. Each set of three coverage panels is followed by two difference heatmaps where UL3 signal has been subtracted. Each row represents a single DAC peak, rank ordered by degree of DAC change in clone ΔH1-A. Top panels show DACs which gained signal (more open chromatin, 8,336), whereas lower panels show DACs which lose ATAC-seq signal (more closed chromatin, 12,285). Metaprofiles are graphed above each heatmap for data separated by direction of accessibility change with a shaded region above and below each metaprofile to indicate one standard error.

**Supplemental Figure 5.**
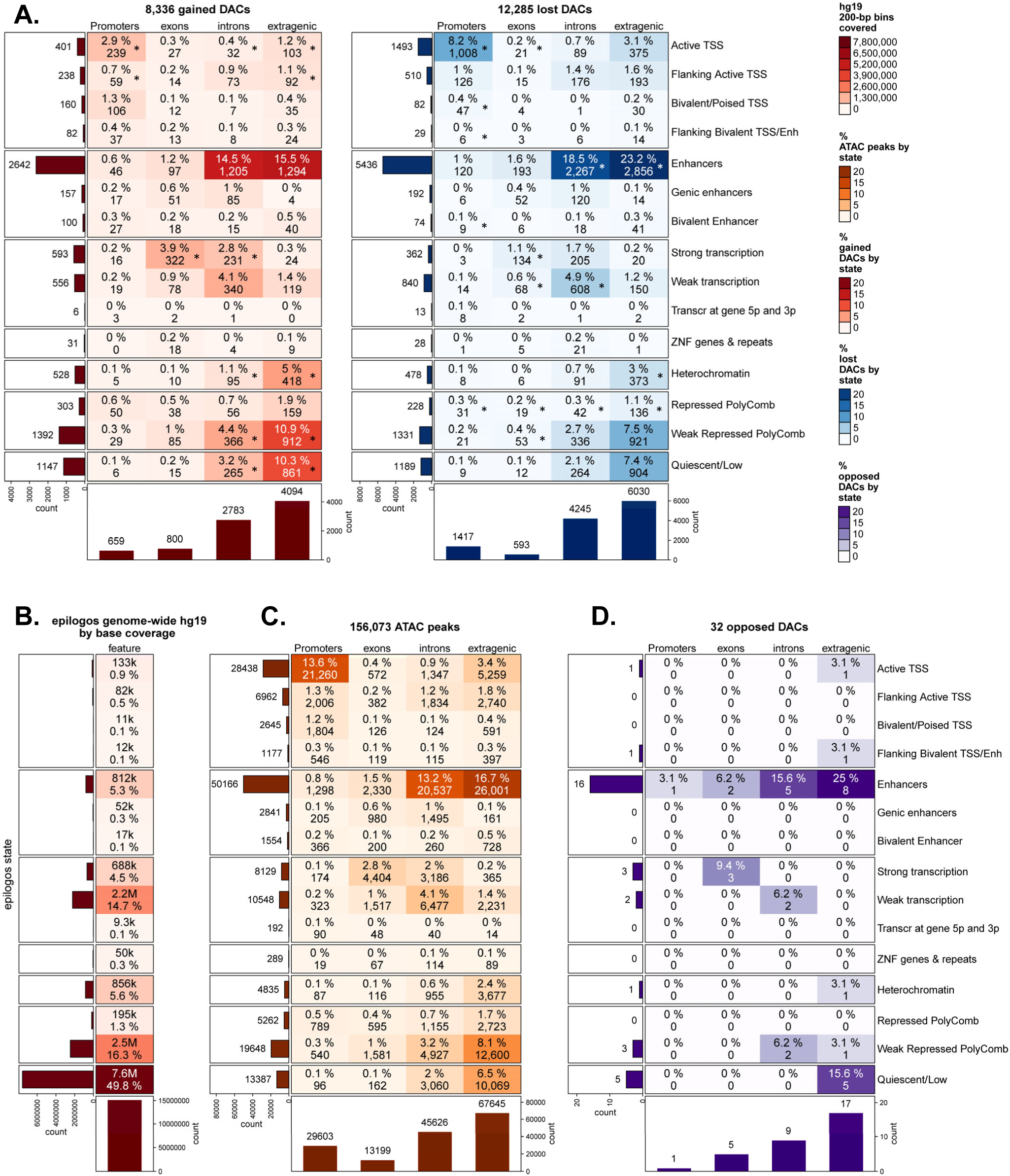
Expanded Epilogos Chromatin and DAC Intersection. **A.** Proportional (%) and count (integer) heatmap of gained DAC locations (left) and lost DAC locations (right) intersected with Epilogos chromatin state (rows) and genic annotation (columns; promoter, exon, intron, extragenic). Bar graphs of row sums (left) and column sums (bottom) shown. Shading was performed based on the proportional representation of the DACs. Asterisks indicate bins for which observed DAC proportions meet statistical significance (p-value <2^-16^) as different relative to all ATAC-seq peaks and Pearson residuals are in Supplemental Table 2. **B.** Heatmap of the relative enrichment of the complete list of Epilogos/ChromHMM chromatin states in the hg19 genome build, broken into 200bp bins. Bin count and relative percentage represented in each chromatin state bin are shown within each cell and each row shows the value for the Epilogos observed chromatin state across 127 cell samples **C.** Proportional (%) and count (integer) heatmap of all called ATAC-seq peak locations intersected with Epilogos chromatin state (rows) and genic annotation (columns). Bar graphs of row sums (left) and column sums (bottom) shown. Shading was performed based on the proportional representation of the DACs. **D.** Heatmap showing the 32 identified DACs which change their accessibility in the H1.4-deficient cells but do so in opposite directions in the two ΔH1.4 CRISPR/Cas9 clones.

**Supplemental Figure 6:**
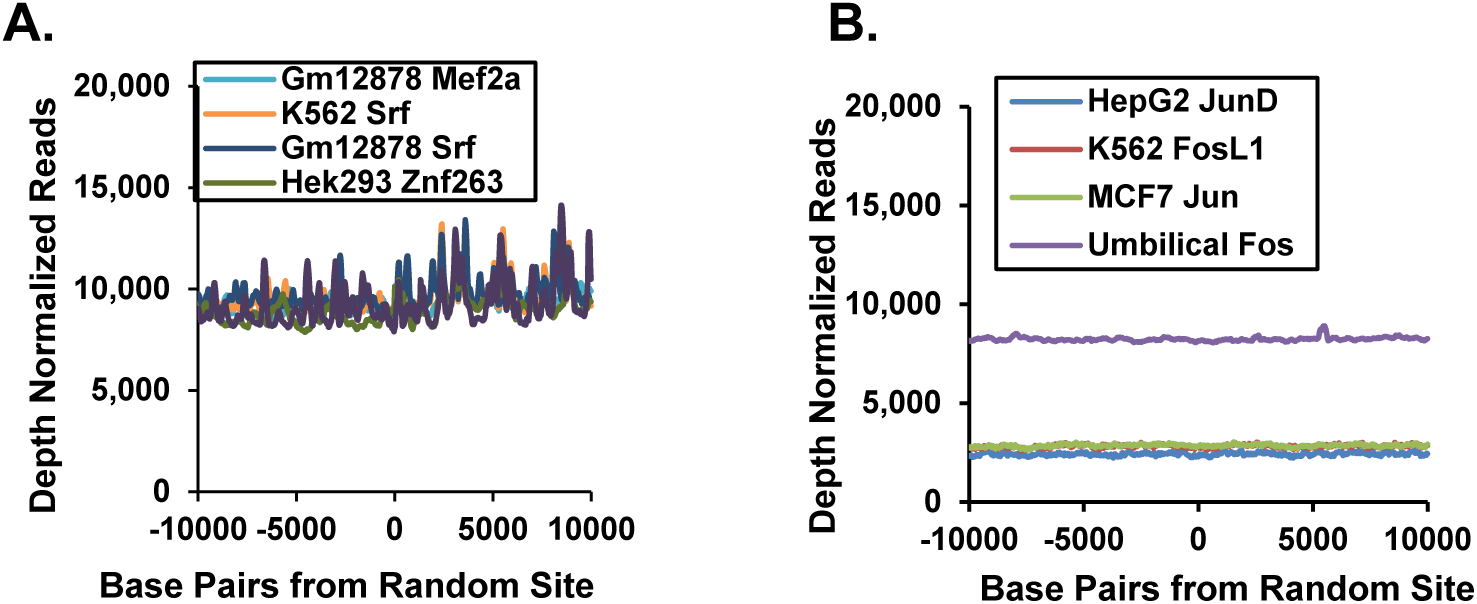
Negative Control Metaplots of Transcription Factor Binding Events. A balanced set of 30,000 random genomic positions were selected and the ENCODE ChIP-seq data for transcription factors identified as motif enriched at the gained DAC peaks (left) or less accessible DAC peaks (right) were s around these sites (+/-10kb) and observed only background signal.

**Supplemental Figure 7:**
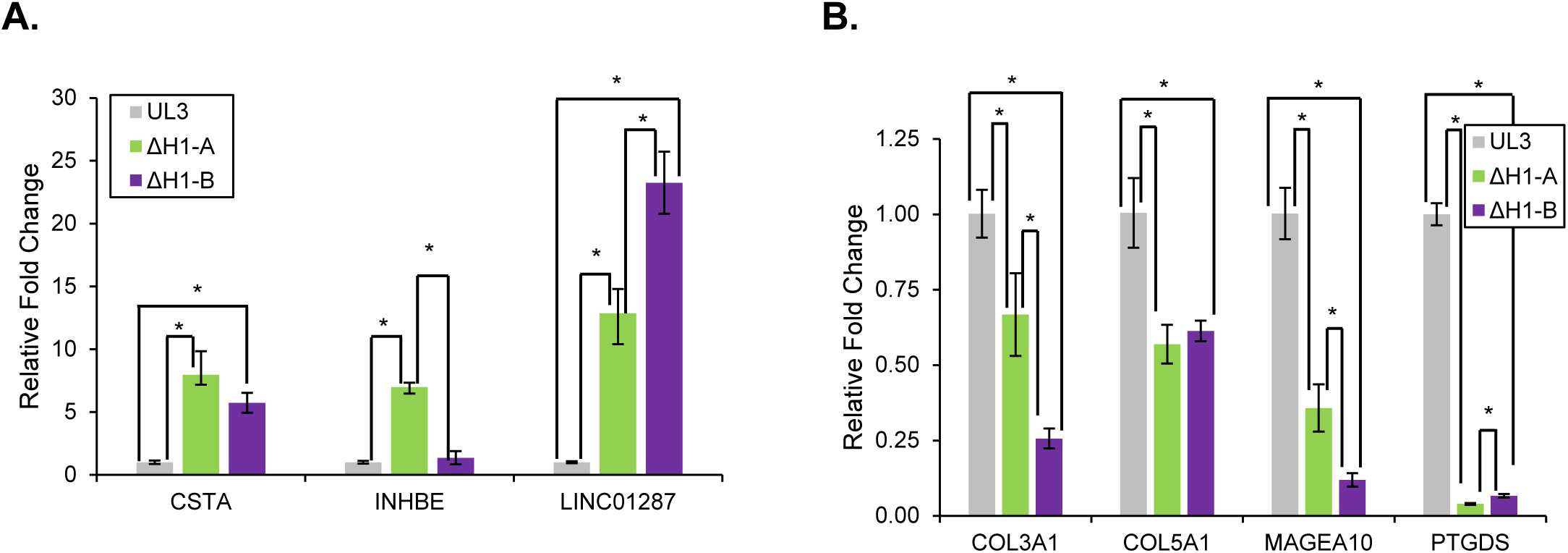
Validation of 4sU-seq findings by RT-qPCR. Validation of identified 4sU-seq DEGs was performed using RT-qPCR in biological quadruplicate for the ΔH1-A (green) and ΔH1-B (purple) ΔH1.4 CRISPR/Cas9 clones relative to the expression in UL3 cells (grey). Both upregulated (**A**) and downregulated genes (**B**) were validated as well as cell-line specific changes (INHBE). Data were normalized to the geometric mean of *POLR3GL*, *RPL13*, *TUBULIN* and *GAPDH* levels, and the and error bars indicate standard deviations with asterisks denoting pval≤0.05.

## Notes

### Competing Interest Statement

The authors have declared no competing interest.

